# The Ebola virus matrix protein clusters phosphatidylserine, a critical step in viral budding

**DOI:** 10.1101/2021.06.08.447555

**Authors:** Monica L. Husby, Souad Amiar, Laura I. Prugar, Emily A. David, Caroline B. Plescia, Kathleen E. Huie, Jennifer M. Brannan, John M. Dye, Elsje Pienaar, Robert V. Stahelin

**Author notes:** Corresponding author (RVS). These authors contributed equally to this work (Souad Amiar, Laura I. Prugar). These authors also contributed equally to this work (Emily A. David, Caroline B. Plescia, Kathleen E. Huie).

## Abstract

Phosphatidylserine (PS) has been shown to be a critical lipid factor in the assembly and spread of numerous lipid enveloped viruses. Here, we describe the ability of the Ebola virus (EBOV) matrix protein eVP40 to induce clustering of PS and promote viral budding *in vitro*, as well as the ability of an FDA approved drug, fendiline, to reduce PS clustering subsequently reducing virus budding and entry. To gain mechanistic insight into fendiline inhibition of EBOV replication, multiple *in vitro* assays were employed including imaging, viral budding and viral entry assays. Fendiline reduced the PS content in mammalian cells and PS in the plasma membrane, reducing the ability of VP40 to form new virus particles. Further, particles that do form from fendiline treated cells have altered particle morphology and decreased infectivity capacity. These complementary studies reveal the mechanism by which filovirus matrix proteins cluster PS to enhance viral assembly, budding, and spread from the host cell while also laying the groundwork for fundamental drug targeting strategies.

## Introduction

Ebola virus (EBOV), which was first discovered in 1976, has been of much concern since the unprecedented 2014-16 outbreak in Western Africa (Breman *et al*, 2016). These fears were exacerbated with the 2018-2020 EBOV outbreak in the Democratic Republic of Congo (>2200 fatalities) as well as a small outbreak there in 2021. The FDA approved an EBOV vaccine (efficacy when administered prior to virus exposure) in 2019 and a monoclonal antibody cocktail in 2020 (Graul *et al*, 2020; Mullard 2020); however, the duration and breadth of these outbreaks underscore the dangers of reoccurring outbreaks and the imminent need to develop small molecule counter measures to treat patients who test positive for EBOV. Further, there is still a large gap in knowledge in how EBOV hijacks host cell components to replicate and spread from cell-to-cell, elucidation of which may identify new drug targets.

In the Filoviridae family, EBOV and Marburg virus (MARV) are lipid enveloped negative-sense single stranded RNA viruses (Banadyga *et al*, 2016; Mühlberger 2007). One commonly overlooked characteristic of many pathogenic viruses, including EBOV and MARV, is their lipid envelope, which is acquired from the host cell they infect. Furthermore, lipid enveloped negative-strand RNA viruses possess limited viral machinery, often encoding for just a handful of viral proteins. Amongst these viral proteins is the multi-functional matrix protein. These matrix proteins, including the VP40 protein of EBOV (eVP40) and MARV (mVP40), are essential to viral assembly and egress (Stahelin 2014; Madara *et al,* 2015). In fact, independent expression of eVP40 or mVP40 leads to the production of virus-like particles (VLPs), nearly indistinguishable from infectious virions (Jasenosky *et al*, 2001; Noda *et al*, 2002; Licata *et al*, 2003). Although these matrix proteins travel through different trafficking pathways within cells, they coalesce at the plasma membrane (PM) to form the viral matrix, which directs viral assembly, budding and the acquisition of their characteristic lipid envelope (Panchal *et al*, 2003; Jasenosky & Kawoaka 2004; Wang *et al*, 2010; Bornholdt *et al*, 2013; Oda *et al*, 2015; Amiar & Stahelin 2020; Wan *et al*, 2020). Importantly, phosphatidylserine (PS) has been implicated in recruiting matrix proteins to the PM and coordinating the assembly of progeny virions (Adu-Gyamfi *et al*, 2015; Wijesinghe & Stahelin 2015; Bobone *et al*, 2017; Del Vecchio *et al*, 2018).

While lipids play a critical role in assembly of new viral particles, lipids are also actively involved in viral entry in a phenomenon known as “apoptotic mimicry”. Apoptotic mimicry is central to the efficient entry of numerous lipid-enveloped viruses (Jemielity *et al*, 2013; Moller-Tank *et al*, 2013; Carnec *et al*, 2015). During apoptotic mimicry, PS is transferred from the inner to the outer leaflet of the PM; this causes PS to become a component of the outer viral envelope during infection (Adu-Gyamfi *et al*, 2015; Amara & Mercer 2015; Nanbo & Kawoaka 2010). Subsequently, the exposed PS in the viral envelope is recognized by target cell receptors for viral uptake, continuing the viral lifecycle (Kondratowicz *et al*, 2011; Moller-Tank *et al*, 2013; Brunton *et al*, 2019).

The two bilayers of the PM have varying compositions of four main phospholipid classes asymmetrically distributed across the two bilayers (Bell *et al*, 1981; Van Meer *et al*, 2008). However, the most abundant anionic lipid within the inner leaflet of the PM is PS, a frequent participant in peripheral protein recruitment (Cho & Stahelin 2005; Stahelin *et al*, 2014; Kerr *et al*, 2018). Extensive work has looked at the dynamic nature of lipids within the PM, including PS, and their tendency to cluster into domains several hundred nanometers in size (Fairn *et al*, 2011; Hirama *et al*, 2017; Bobone *et al*, 2017). Clustering of anionic lipids into domains enriches regions of the PM with anionic charge, creating a platform for electrostatic interactions at the PM interface for peripheral protein recruitment. This phenomenon has been reported between PS and the matrix protein of influenza A virus (Bobone *et al*, 2017). Although significant work has underscored the importance of PS in filovirus budding and entry (Moller-Tank *et al*, 2013; Moller-Tank & Maury 2014; Soni & Stahelin 2014; Stahelin 2014; Adu-Gyamfi *et al*, 2015; Del Vecchio *et al*, 2018; Brunton *et al*, 2019), the molecular details of the interaction has not been explored in the context of the lateral organization of PS, matrix assembly or implications on viral spread.

The FDA approved drug, fendiline, has been reported to reduce PS levels within the PM inner leaflet (Cho *et al*, 2016; van der Hoeven *et al*, 2018), which was sufficient to inhibit the oncogenic protein K-Ras PM localization and signaling (van der Hoeven *et al*, 2013; van der Hoeven *et al*, 2018). Fendiline was initially approved by the FDA in the 1970s as a non-selective calcium channel blocker to treat coronary heart disease (Bayer *et al*, 1987); however, these recently identified off target properties were found to be calcium independent and associated with the indirect inhibition of acid sphingomyelinase (ASM) (Cho *et al*, 2016; van der Hoeven *et al*, 2018). eVP40 has been shown to utilize PS for PM localization, assembly, and production of progeny virions (Adu-Gyamfi *et al*, 2015; Del Vecchio *et al*, 2018; Johnson *et al*, 2021); however, detailed molecular insight into this relationship is lacking. To delineate the molecular architecture and requirements of PS concentration on VP40 assembly, oligomerization and budding, we employed biochemical and biophysical assays *in vitro* and in cells. We also tested the ability and mechanism by which eVP40 clusters PS *in vitro* and in cells. We hypothesized that reduction of PS from the PM with fendiline treatment would perturb EBOV assembly and inhibit viral budding. Lastly, fendiline treatment was tested as a potential therapy for inhibition of EBOV budding and spread in biosafety level (BSL)-2 and BSL-4 models of infection.

## Results

### EBOV VP40 localizes to PS enriched regions in synthetic membranes and in living cells

eVP40 has been reported to have selectivity and high affinity for PS (Ruigrok *et al*, 2000; Scianimanico *et al*, 2000; Adu-Gyamfi *et al*, 2015; Del Vecchio *et al*, 2018) and PI(4,5)P_2_ (Johnson *et al*, 2016; Johnson *et al*, 2018), both *in vitro* and in cells. To establish an *in vitro* system for eVP40-PS colocalization, we employed a giant unilamellar vesicle (GUV) system with fluorescently labelled His_6_-eVP40 and a fluorescent PS molecule, TopFluor® TMR-PS (tetramethylrhodamine-PS (TMR-PS)). Dipalmitoylphosphatidylserine (DPPS) was chosen for this assay as eVP40 associates with saturated PS containing membranes *in vitro* more significantly than PS mono- or diunsaturated containing membranes (data not shown). In GUVs supplemented with 40% DPPS, a homogeneous ring structure of eVP40-Alexa488 surrounding the GUV membrane (Fig EV1A, *middle panel*) was observed and a random plot profile analysis indicated a small overlap of the two fluorescent signals at the membrane (Fig EV1C, indicated by the asterisk). To investigate further the ability of eVP40-Alexa488 to associate with membranes similar to the PM, GUVs containing both 40% DPPS and 2.5% PI(4,5)P_2_ were used (Fig EV1A *right panel*)(See Material and Methods section for important temperature preparation guidelines). The plot profile analysis of the image in Expanded View Fig 1A revealed a strong overlap between the two fluorescence signals (eVP40 and TMR-PS) (Fig EV1D). The protein enrichment analysis also supported this previous observation with an enrichment index of ∼5.4 ± 2, four times more than membranes devoid of PI(4,5)P_2_ (Fig EV1E). These data are in agreement with the previously published findings that eVP40 requires both PS and PI(4,5)P_2_ for efficient membrane binding and oligomerization and suggest eVP40 is able to enrich PS at sites of VP40 oligomerization.

**Figure 1.**
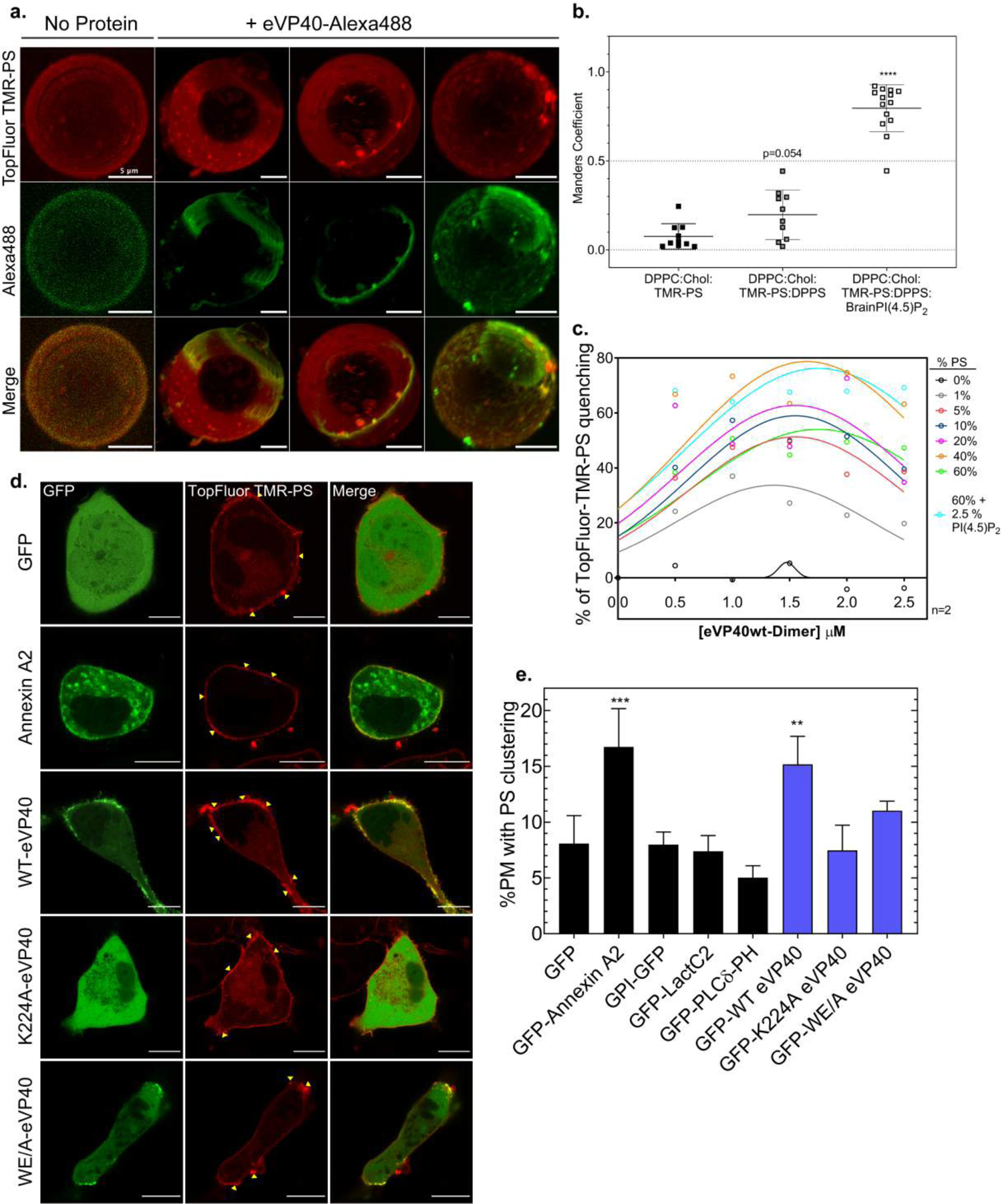
Clustering of PS by eVP40 *in vitro* and in HEK 293 cells. **A** Representative 3D reconstructed confocal images of immobilized GUVs (DPPC:Cholesterol:DPPS:PI(4,5)P_2_:TopFluor® TMR-PS(red)). Left panel: GUVs incubated without eVP40-Alexa488. Right three panels: GUVs incubated with 1.25 µM eVP40-Alexa488 (green). **B** Index of correlation (Mander’s coefficient) between TopFluor® TMR-PS and eVP40-Alexa488 of different GUVs compositions incubated with 1.25 µM eVP40-Alexa488. Values are reported as mean ± s.d. A one-way ANOVA with multiple comparisons was performed (n = 3); ****p<0.0001. **C** % TopFluor® TMR-PS quenching by eVP40 using GUVs (DPPC:Cholesterol:TopFluor®TMR-PS + increasing mol% of PS), 2.5% PI(4,5)P_2_ was added to GUVs with 60% PS. Fluorescence spectra were recorded (Ex: 547 nm; Em: 550-600 nm); n=2. **D** Representative confocal images of HEK293 cells expressing various GFP**-**fused proteins (green) and supplemented with TopFluor® TMR-PS (red); scale bar= 10 µm. Yellow arrows indicate high intensity PS fluorescence regions **E** %PM with PS clusters = area of high intensity fluorescent PS clusters over total plasma membrane area from images in panel D. Black bars are control proteins and blue bars are eVP40 proteins. Values are reported as mean ± s.d.; N>18, n=3; A one-way ANOVA was performed with multiple comparisons compared to the control GFP %PS clustering (***p=0.0007, **p=0.004). DPPC: dipalmitoyl-phosphatidylcholine; DPPS: dipalmitoyl-phosphatidylserine; GUVs: giant unilamellar vesicles; PS: phosphatidylserine; PM: plasma membrane.

To expand upon our findings, we investigated if eVP40 localized to PS enriched regions within the plasma membrane of live cells. To visualize PS and protein localization simultaneously, we transiently expressed EGFP-fused proteins in HEK293 cells and supplemented the cells with TMR-PS immediately prior to imaging (Fig EV2A). To test the hypothesis that eVP40 localizes to PS enriched regions of the PM, we examined the plot profile analysis of TMR-PS and eVP40 by expressing functionally unique EGFP fused proteins: WT-eVP40, Lact-C2 (a PS binding reporter), K224A-eVP40 (a PS-binding residue mutant (Del Vecchio *et al*, 2018), and WE/A-eVP40 (oligomerization deficient mutant (Adu-Gyamfi *et al*, 2012; Bornholdt *et al*, 2013). The fluorescence profile of EGFP-WT-eVP40 vs. TMR-PS revealed a strong overlap between the two fluorophores (Fig EV2A,D). This cellular data corroborates our *in vitro* data, demonstrating that EGFP-eVP40 localizes to PS enriched regions of both model membranes and in the PM of cells. Additionally, there was no significant fluorescence signal overlap between the EGFP-K224A-eVP40 mutant and TMR-PS (Fig EV2A,E), which supports the requirement of PS binding for PM localization of eVP40 (Del Vecchio *et al*, 2018). Importantly, random plot profile analysis revealed a moderate overlap in the fluorescence signals of the oligomerization deficient mutant WE/A-eVP40 and TMR-PS (Fig EV2A,F). This is important to note as this protein is still able to interact with PS at the PM, however, is unable to properly oligomerize (Hoenen *et al*, 2010; Adu-Gyamfi *et al*, 2012; Adu-Gyamfi *et al*, 2013). These results suggest VP40 interacts with PS at the PM inner leaflet as a dimer without significant oligomerization, in line with VP40 *in vitro* lipid-binding (Del Vecchio *et al*, 2018).

### EBOV-VP40 enhances clustering of PS in synthetic membranes and in living cells

Biophysical and molecular studies into PS dynamics in model membranes and living cells revealed that PS basally distributes into clustered domains enriched with PS (Fairn *et al*, 2011; Hirama *et al*; 2017; Bobone *et al*, 2017). Interestingly, cellular proteins such as Annexins are known to significantly enhance the clustering of PS (Menke *et al*, 2005) and viral proteins such as M1 of Influenza A virus have a selectivity for these PS clusters (Bobone *et al*, 2017). However, detailed examination of PS clustering and whether filovirus matrix proteins such as eVP40 alter the organization of PS has not yet been explored.

Confocal 3D reconstruction of GUVs (with DPPS and PI(4,5)P_2_) indicated different structures of TMR-PS clusters observed where eVP40-Alexa488 fluorescence was enriched (Fig EV1A, *three right columns*). Furthermore, the Mander’s coefficient index of correlation was quantified (between the TMR-PS and eVP40-Alexa488) with varying lipid compositions (Fig 1B) demonstrating a statistically significant increase in PS clustering when eVP40-Alexa488 was incubated with GUVs containing both DPPS and PI(4,5)P_2_ (Fig. 1b; *p<0.0001). These results suggest that eVP40 induced PS clustering *in vitro,* which was significantly enhanced in the presence of both PS and PI(4,5)P_2_, akin to the lipid composition typically found in the PM inner leaflet.

Next, we were interested if eVP40 was able to induce PS clustering in synthetic membranes in the absence of PI(4,5)P_2_. Therefore, we performed a TMR self-quenching experiment as described previously (Zhao *et al*, 2010; Wen *et al*, 2018). The TMR fluorescent group has potent self-quenching properties when two molecules or more are brought in close distance from each other. We tested the ability of eVP40 to undergo TMR self-quenching when fluorescent PS was incorporated in lipid vesicles, as a secondary effect to eVP40-induced PS-clustering. We also tested different eVP40 concentrations and different PS ratios to investigate if TMR self-quenching was concentration dependent. The strongest TMR self-quenching in all membranes was observed at 1.5 µM eVP40 (Fig 1C). As expected, higher PS molar ratios resulted in stronger eVP40 induction of TMR self-quenching except for a high molar ratio of DPPS (60% molar ratio). However, the addition of PI(4,5)P_2_ at 2.5% molar ratio to 60% DPPS-containing membranes rescued TMR self-quenching similarly to 40% DPPS-containing membranes. Further, DPPS-containing membranes at 60% molar ratio with or without PI(4,5)P_2_ displayed a maximum of TMR-PS self-quenching at 2 µM eVP40. These observations may indicate a saturation of the liposome membranes with eVP40 at high PS concentrations. Altogether, this assay demonstrated that eVP40 is able to cluster PS, which is enhanced in the presence of PI(4,5)P_2_.

We next examined if eVP40 enhanced PS clustering in the PM of cells. As previously mentioned, PS selectively localizes into clustered regions (Fairn *et al*, 2011; Hirama *et al*; 2017; Bobone *et al*, 2017); and we were able to detect a basal level of PS enriched clusters in our control GFP expressing cells, with PS clusters accounting for approximately ∼8% of the PM (Fig 1D *top panel* and Fig 1E). Our method was further validated by expressing an additional control protein with a glycosylphosphatidylinositol membrane anchor conjugated to GFP (GFP-GPI), which revealed PS clusters in ∼8% of the PM area (Fig 1E; Representative image in Fig EV3B *top panel*).

We next sought to determine if our technique accurately captured enhanced PS clustering using the established EGFP-Annexin A2 PS reporter (Menke *et al*, 2005). As shown in Fig 1D-E, expression of EGFP-Annexin A2 significantly enhanced PS clustering roughly 2-fold, compared to EGFP expressing cells (***p=0.0001). Taken together, these findings corroborate the previously reported effect of Annexin A2 on PS organization (Menke *et al*, 2005), as well as validate the method developed for our assay. In contrast, expression of EGFP-PLCδ-PH (PI(4,5)P_2_) or EGFP-LactC2 did not significantly alter the extent of PS clustering (Fig 1E). This confirmed that transient expression of fluorescently conjugated lipid-binding proteins is not sufficient to enhance PS clustering at the PM.

Next, we evaluated the effect of eVP40 expression on PS organization across the PM. EGFP-WT-eVP40 increased PS clustering by ∼2 fold (*p=0.004), similar to the PS clustering observed with Annexin A2 (Fig 1D,E). However, expression of the PS-binding deficient mutant EGFP-K224A-eVP40 showed no significant change in PS clustering (Fig 1D,E), supporting the hypothesis that eVP40 must interact with PS to promote its clustering at the PM. Additionally, to investigate if eVP40 matrix oligomerization was important for PS clustering, we expressed EGFP-WE/A-eVP40 in HEK293 cells. It is important to note that this mutant still colocalizes with PS at the PM (Fig EV2A,F) albeit to a lesser extent than WT (Adu-Gyamfi *et al*, 2012). Although the WE/A-eVP40 and PS interaction is maintained in cells, no significant increase in PS clustering was observed (Fig 1D,E). To the best of our knowledge, this is the first account of a filovirus matrix protein modulating the organization of PS within the PM. Moreover, these results demonstrate that both membrane binding and oligomerization of eVP40 is central to eVP40-mediated PS clustering.

### eVP40 membrane binding and oligomerization are dependent on phosphatidylserine content in lipid membranes

Next, we hypothesized eVP40 may require PS clustering for productive interactions at the PM during assembly. Enrichment of PS within regions of the PM would provide additional PS molecules available to recruit eVP40 to platforms of viral budding. To investigate how increasing the amount of PS within membranes dictates eVP40 membrane affinity, surface plasmon resonance (SPR) was performed with His_6_-eVP40 and large unilamellar vesicles (LUVs) with increasing concentrations of PS (POPC matrix with PS added from 1% to 22 mol% PS; Fig 2A-C). eVP40 displayed moderate binding to LUVs with 1% PS, with an apparent affinity of 2.5 µM (Fig 2A). However, increasing the concentration of PS to 11% increased the apparent affinity of eVP40 to 0.65 µM (Fig 2B). eVP40 displayed even stronger affinity to vesicles with 22% PS, with an apparent affinity (K_d_) of ∼0.18 µM (Fig 2C). These results indicate that by increasing the amount of PS in membranes, the affinity of eVP40 to lipid membranes can be modulated. This finding supports the hypothesis that PS clustering may be a mechanism for the virus to provide the necessary electrostatic contacts needed for matrix assembly during viral production. Once at the PM, VP40 oligomerizes into the extensive matrix that gives rise to the stability and structure of the virion. Previously, Adu-Gyamfi et al (Adu-Gyamfi *et al*, 2015) highlighted the importance of PS in this process, where a cell line deficient in PS synthesis showed a significant reduction in eVP40 oligomerization. Moreover, our confocal clustering data (Fig 1D,E) revealed that eVP40 oligomerization is crucial for modulating PS organization into clustered domains. To investigate how increasing PS concentration alters eVP40 oligomerization, we utilized chemical crosslinking of His_6_-eVP40 which had been incubated with LUVs of increasing PS concentration (Fig 2D,E). Introduction of 15% PS into LUVs led to extensive eVP40 oligomerization beyond dimeric eVP40 compared to when 0% PS LUVs were used (Fig 2D *lane 2*, Fig 1E). We next tested LUVs containing 30% and 60% PS and found that eVP40 oligomerization was even more significantly detected than when just 15% PS was used (Fig 2D *lane 3 and lane 4, respectively*, Fig 1E). Compared to LUVs with 0% PS, both 30% and 60% PS led to a significant increase in eVP40 oligomerization (*p=0.021 and *p=0.017, respectively). Further, eVP40 oligomerization appeared to saturate when 30% PS was included, as increasing PS content to 60% did not increase eVP40 oligomerization (compared to 30% PS). Taken together, these studies suggest a dynamic relationship between PS clustering and eVP40 affinity and oligomerization as a critical step in eVP40 assembly.

**Figure 2.**
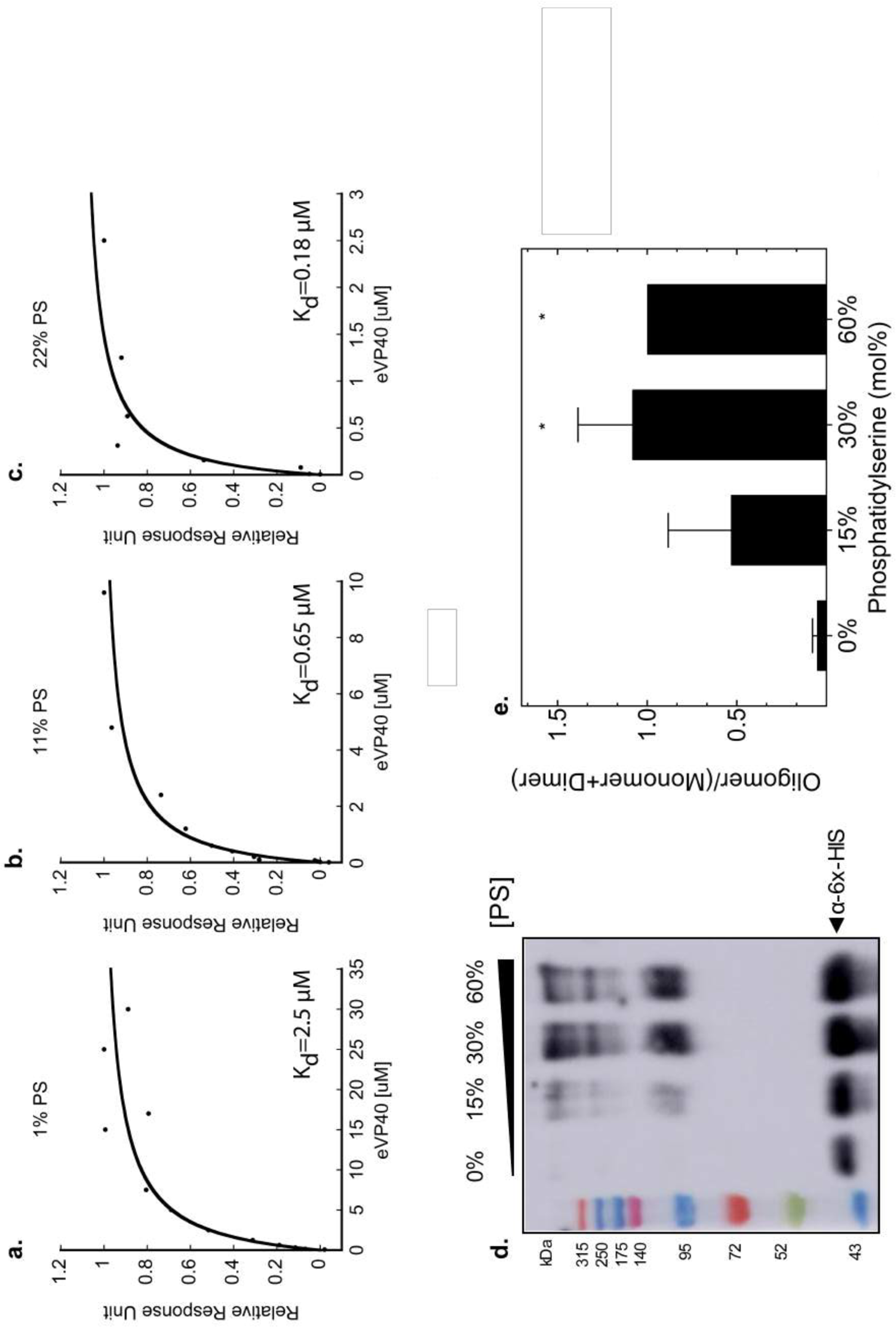
Effect of PS concentration on eVP40 binding affinity to and oligomerization on membranes. **A-C** SPR demonstrates that eVP40 affinity to LUVs increases in relation to PS concentration. **A** Representative normalized sensorgram of His_6_-eVP40 binding to LUVs containing 1% PS indicating an apparent affinity of 2.5 µM. **B** Representative normalized sensorgram of His_6_-eVP40 binding to LUVs containing 11% PS indicating an apparent affinity of 0.65 µM. **C** Representative normalized sensorgram of His_6_-eVP40 binding to LUVs containing 22% PS indicating an apparent affinity of 0.18 µM. **D-E** PS concentration in LUVs enhances the ability of His_6_-eVP40 to oligomerize on membranes. **D** Representative western blot of chemical crosslinking performed on His_6_-WT-eVP40 following incubation with LUVs of varying PS content (detected by Mouse α-His antibody & HRP-Sheep α-Mouse). **E** Oligomerization capacity was determined from the western blot band density ratio of oligomers/(monomer + dimer) from chemical crosslinking experiments. A one-way ANOVA was performed with multiple comparisons compared to the control 0% PS LUVs control (30% PS *p= 0.021; 60% PS *p=0.017). n=3. Values are reported as mean ± s.d.; SPR: surface plasmon resonance; LUVs: large unilamellar vesicles; PS: phosphatidylserine; HRP: horseradish peroxidase.

### Total cellular and plasma membrane levels of phosphatidylserine are reduced by fendiline treatment

A recent study reported an FDA-approved drug, fendiline, inhibited K-Ras PM localization and signaling (Cho *et al*, 2016) and reduced PM PS content in MDCK cells (Cho *et al*, 2016). Therefore, it was our goal to determine if fendiline could also reduce PS levels in the human cell line HEK293, a cell line commonly used in BSL-2 filovirus studies. The initial finding that fendiline reduced PS levels within the PM (40% reduction, IC_50_ ∼3uM) was conducted in MDCK cells using thin-layer chromatography (Cho *et al*, 2016), therefore it had not been established if this effect was cell-type specific. To address this, we first established fendiline’s toxicity in HEK293 cells after 24 and 48 hours of treatment. No significant toxicity was observed in treatments up to 5 µM fendiline (Fig EV4A). To evaluate the effect of fendiline on PS in HEK293 cells, cells were treated with fendiline for 48 hours and lipids were extracted and quantified by liquid chromatography-tandem mass spectrometry (LC-MS/MS). We observed a significant reduction in cellular PS levels compared to DMSO treated cells, after 48 hour treatment with 1 µM fendiline (∼18% reduction; *p=0.012) and 5 µM fendiline (∼30% reduction; ***p=0.0003) (Fig 3A). Fendiline exhibited no selectivity in reducing different PS species, as 5 µM fendiline reduced long chain (C>38) and saturated PS species nearly equally (Fig EV4B). It is important to note that the effect of fendiline on PS was specific in that fendiline treatment had no significant effect on another anionic phospholipid, phosphatidic acid (Fig EV4C). Therefore, our data supports the reported finding that fendiline reduced total cellular levels of PS, and that the effect is not cell dependent.

**Figure 3.**
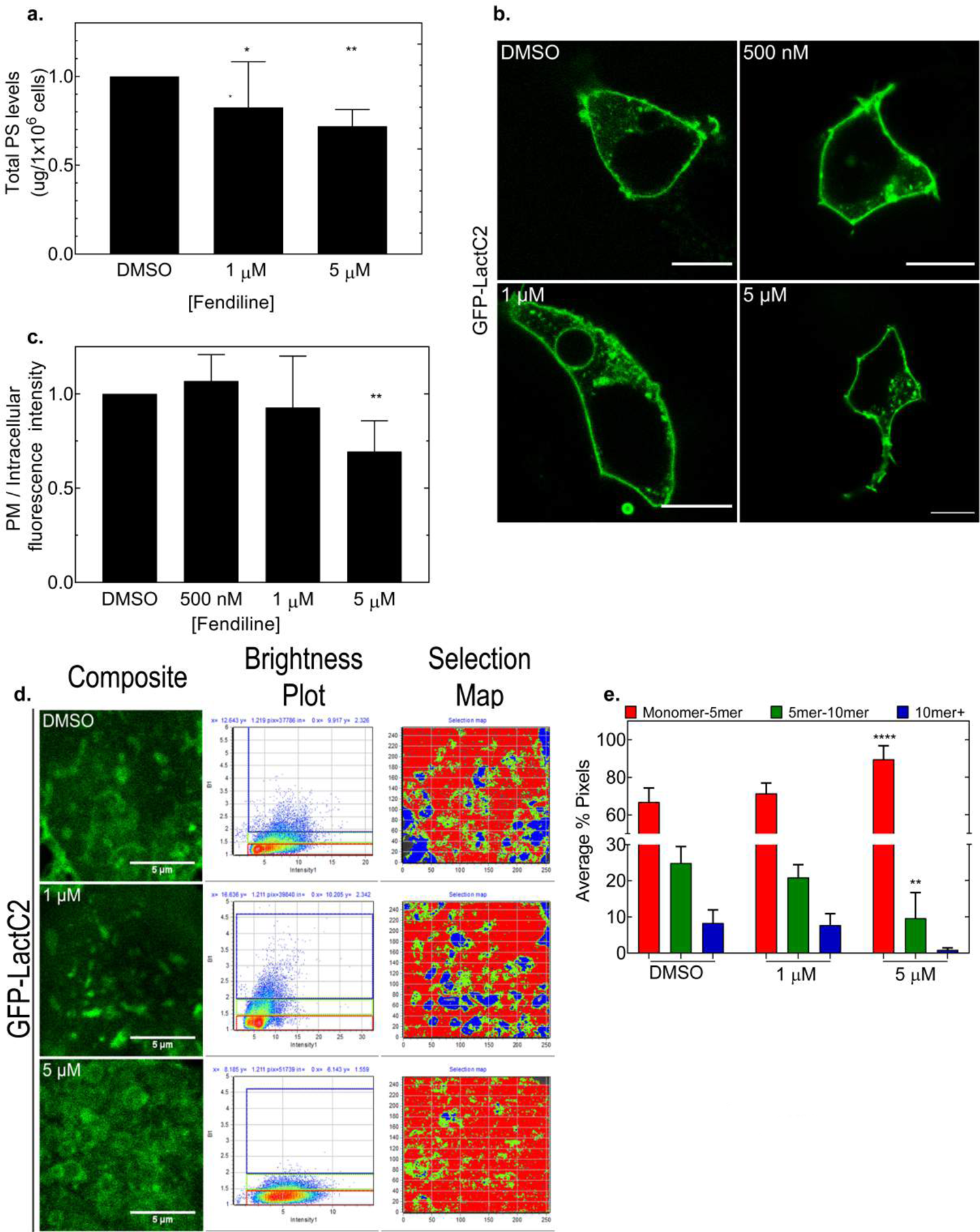
PS concentration, localization, and dynamics in fendiline treated HEK293 cells. **A** Lipidomic analysis (LC/MS/MS) of total lipids extracted from HEK293 cells treated with the indicated concentration of fendiline (48 hours) demonstrated a significant reduction of total cellular PS levels. Values are normalized to DMSO control and are reported as mean ± s.d.; n=3; A one-way ANOVA was performed with multiple comparisons to the control DMSO (*p=0.0120, ***p=0.0003). **B-C** Analysis of PS plasma membrane localization in response to fendiline treatment in HEK293 cells. **B** Representative confocal images from live cell imaging of HEK293 cells expressing GFP-LactC2 and treated with fendiline for 48 hours; scale bars= 10 µm. **C** Effect of fendiline on PS plasma membrane localization was calculated by the ratio of GFP fluorescence at the plasma membrane intensity/intracellular intensity. Values are normalized to DMSO control and are reported as mean ± s.d.; N>15, n=3; A one-way ANOVA was performed with multiple comparisons compared to the DMSO control (**p=0.0031) **D-E** Analysis of PS clustering in HEK293 cells in response to fendiline treatment through N&B analysis. **D** *Left panel:* Representative images from time-lapse (30 frames) imaging of HEK293 expressing GFP-LactC2 and treated with fendiline for 48 hours; scale bar= 5 µm. *Middle panel:* Brightness and Intensity plots for each representative image. *Right panel:* Selection map correlating each pixel in the representative image to an oligomerization state (b value) (red: monomer-5mer, green: 5mer-10mer, blue: >10mer). **E** Average % pixels quantification from panel (d)= Percentage of GFP-LactC2 with brightness values corresponding to monomer-5mer (∼1.-1.5), 5mer-10mer (∼1.5-1.9) and >10mer (>1.9) over the total pixels within each image. Values are reported as mean ± s.d.; N≥9, n=3; A two-way ANOVA was performed with Dunnett’s multiple comparisons comparted to the control DMSO % average pixels (****p<0.0001, **p=0.0043).GFP-LactC2: phosphatidylserine sensor; N&B: Number & Brightness analysis; PM: plasma membrane; PS: phosphatidylserine.

As PS is an integral anionic component of the PM inner leaflet, we sought to confirm that fendiline treatment also reduced PS levels within the PM in HEK293 cells. PS localization within the PM has been readily studied by expressing EGFP-LactC2 in mammalian cells (Yeung *et al*, 2008; Kay *et al*, 2012). Therefore, HEK293 cells expressing EGFP-LactC2 were imaged at 24 hours (Fig EV4D) and 48 hours (Fig 3B) post-treatment with increasing concentrations of fendiline. Single doses of 500 nM fendiline had no effect on EGFP-LactC2 PM localization at 24 or 48 hours post treatment (Fig EV4E and Fig 3C, respectively). However, we found a ∼30% reduction in PM EGFP-LactC2 localization after 24 hours of treatment for both 1 µM (**p=0.0003) and 5 µM fendiline (**p=0.0045) (Fig EV4E). However, a single dose of 1 µM fendiline treatment did not significantly affect Lact-C2 PM localization after 48 hours of treatment (Fig 3C). Conversely, a single dose of 5 µM fendiline significantly reduced Lact-C2 PM localization even at 48 hours post treatment (∼30% reduction; **p=0.0031; Fig. 3c), a reduction similar to that observed at 24 hours post treatment.

Fendiline has been known to target L-type calcium channels in the T-tubules (invaginations of the plasma membrane) of muscle cells thereby inhibiting calcium movement and acting as a vasodilator (Bayer & Mannhold 1987). However, in a number of non-muscle cell lines, fendiline has been shown to increase calcium levels, including hepatoma cells (Cheng *et al*, 2001), oral cancer cells (Huang *et al*, 2009), PC prostate cancer cells (Jan *et al*, 2001), and canine kidney cells (MDCK) (Jan *et al*, 2000). Fendiline was also specific in the K-Ras inhibitory affect among a number of L-type calcium channel blockers (van der Hoeven *et al*, 2013) strongly suggesting the fendiline mechanism of action is directly linked to PS depletion and not changes in calcium. Fendiline was shown to inhibit ASM (van der Hoeven *et al*, 2015) thereby reducing plasma membrane sphingomyelin, ceramide, PS and cholesterol. Restoration of ASM activity lead to the relocalization of the PS reporter LactC2 to the PM (van der Hoeven et al, 2015). Lastly, studies in HEK293 cells, the major cell line used herein, demonstrated endogenous calcium channel currents could be distinguished from heterologous expression of L-type calcium channels (Berjukow *et al*, 1996).

Thus, based upon rigor of prior research the most likely hypothesis is that fendiline inhibits ASM to alter sphingomyelin levels, which are necessary to maintain proper PS levels in the plasma membrane. Recently, fendiline analogs have been developed with nanomolar potency and have been shown to displace ASM, which is membrane bound in acidic lysosomes leading to ASM degradation (Wang *et al*, 2021) While the direct mechanism of PS depletion in the PM is not fully understood, transbilayer communication may take place across the plasma membrane between sphingomyelin levels on the outer leaflet and phosphatidylserine and cholesterol on the inner leaflet (Llorente *et al*, 2013; Skotland & Sandvig 2019).

### Fendiline reduces PS clustering

Next, we hypothesized that reduced levels of PS within the PM would therefore reduce the degree of PS clustering. To determine if fendiline treatment reduced the degree of PS clustering, we utilized the Number & Brightness technique (N&B). N&B is a quantitative fluorescence microscopy technique that allows one to detect the aggregation state of proteins with pixel resolution in real time (Digman *et al*, 2008). Previously, N&B was used to quantify PS clustering by analyzing the N&B profile of EGFP-LactC2 (Bobone *et al*, 2017). To accurately capture PS clustering at the PM, imaging was performed at a focal plane near the cell surface (Fig EV4F).

To evaluate PS clustering, HEK293 cells expressing EGFP-LactC2 were treated with control or fendiline for 48 hours and the EGFP-LactC2 N&B profile was examined (Fig 3D,E). Three different cluster bin sizes were examined, 1—5, 5-10 and >10. The average percentage of pixels in each bin was calculated and plotted (Fig 3E). Within control-treated cells, significant aggregation of EGFP-LactC2 was observed, with 25% present in complexes of 5-10 LactC2 molecules and ∼10% in complexes of >10 LactC2 molecules (Fig 3D *top panel* and Fig 3E). This corroborates previous work investigating PS clustering, which found EGFP-LactC2 clusters up to 15 molecules in size (Bobone *et al*, 2017). Treatment of cells with 5 µM fendiline abolished the presence of EGFP-LactC2 complexes (>10 molecules) and significantly reduced the number of complexes of 5-10 LactC2 molecules large (from ∼8% in DMSO to ∼0% in 5 µM fendiline; **p=0.0043) (Fig 3D *bottom panel* and Fig 3E). Moreover, there was a significant (∼23%) increase in EGFP-LactC2 complexes of ∼1-5 molecules in size in cells treated with 5 µM fendiline compared to control treated cells (****p<0.0001) (Fig 3D *bottom panel* and Fig 3E). Taken together, this data suggests that fendiline treatment disrupted large PS-dependent LactC2 complexes which was compensated by an increase in smaller PS-dependent complexes. Therefore, fendiline may possess antiviral properties by disassembling PS enriched regions that would otherwise have been used as platforms for viral assembly.

While the mechanism by which fendiline reduces PS clustering is not known, it has been hypothesized that sphingomyelin changes on the outer plasma membrane can alter the distribution of PS in the inner leaflet (Skotland & Sandvig 2019). This is thought to be attributed to the long fatty acid chains of sphingomyelin interacting with the hydrophobic chains of PS, especially that of PS 18:0/18:1 (Skotland & Sandvig 2019), the most predominant PS species in the plasma membrane. The changes in cholesterol in the plasma membrane could also be partially responsible as optimal levels of sphingomyelin are required for maintenance of cholesterol and PS levels in the plasma membrane inner leaflet (Maekawa *et al*, 2016 Sci Rep) and PS is necessary for proper cholesterol distribution across the plasma membrane (Maekawa & Fairn 2015).

### Fendiline significantly inhibits authentic EBOV replication

To determine if the FDA-approved drug fendiline was able to inhibit authentic filovirus replication and spread, we first established the toxicity of fendiline in Vero E6 cells (Fig EV5A) an established model for BSL-4 filovirus studies (Moe *et al*, 1981), and then monitored the efficacy of fendiline at inhibiting EBOV replication in a BSL-4 setting. Vero E6 cells were used to examine filovirus replication 48, 72 and 96-hours post-infection at different multiplicity of infection (MOI) with EBOV (Kikwit) (Fig 4). Following infection, plates were separated into three post-infection treatment groups (day 0, every day dosing (e.d.), or every other day dosing (e.o.d.)).

**Figure 4.**
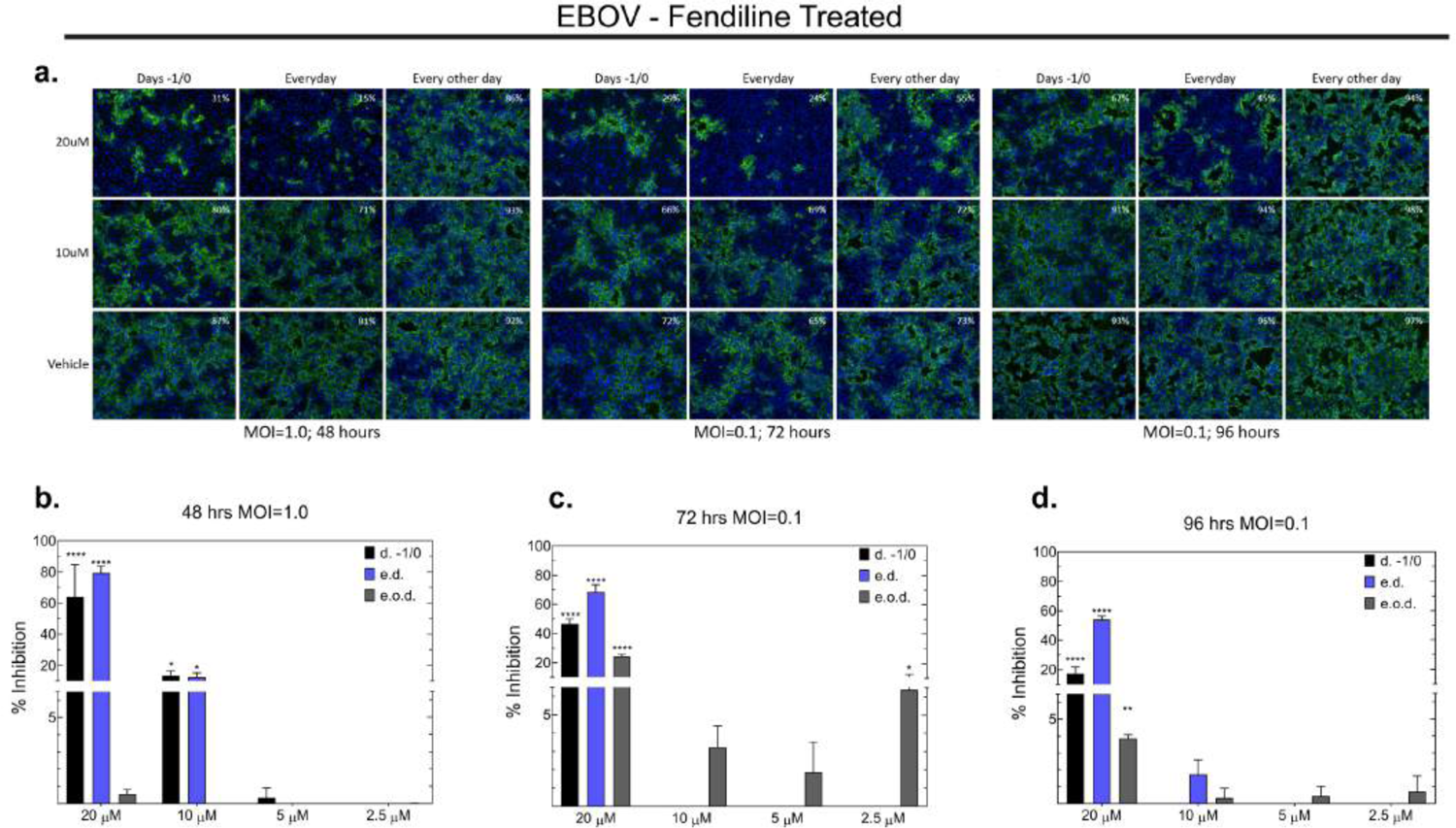
Evaluation of fendiline efficacy in the inhibition of authentic EBOV spread. **A-D** Effect of fendiline on EBOV infection. **A** Representative confocal images of Vero E6 cells infected with EBOV (Kikwit) at the indicated MOI and treated with the indicated concentration of fendiline. Cells were pretreated 24 hours prior to infection with the indicated concentration of fendiline. Post infection, cells were treated 1 hour later (d −1/0), treated every day (e.d), or treated every other day (e.o.d) and fixed at either 48 hours, 72 hours or 96 hours post infection. (green=EBOV; blue= nuclei). White numbering in top right corner indicates %infection. **B-D** Quantification of % inhibition of EBOV by fendiline. **B** 48 hours (MOI 1.0) **C** 72 hours (MOI 0.1) **D** 96 hours (MOI 0.1). Values are reported as mean ± s.d. A one-way ANOVA was performed with multiple comparisons was performed. n=3. EBOV: Ebola virus; MOI: multiplicity of infection; d. −1/0: treatment 1 hour after infection; e.d.: treatment every day; e.o.d.: treatment every other day.

Fendiline was most effective at reducing EBOV infection in vitro at the highest 20 µM concentrations in each treatment group with statistically significant inhibition observed for EBOV at each time point and each treatment group (excluding EBOV 48 hours, e.d.)(****p<0.0001, **p<0.0066) (Fig 4). Percent inhibition was directly affected by timing of treatments following infection. Furthermore, EBOV infection treatment with fendiline e.d. had the highest inhibition on viral spread at each time point for 20 µM fendiline treatments. Cells of the e.o.d. treatments group, which did not receive treatment immediately following infection with virus, had a dramatically reduced degree of inhibition as compared to the day 0 and e.d. treatment groups, both of which received fendiline immediately following viral infection of one hour.

### Fendiline reduced EBOV-VP40 localization to the plasma membrane

As both EBOV and MARV-VP40 assembly at the PM is in part governed by PS, we first analyzed both EGFP-eVP40 and EGFP-mVP40 PM localization in cells treated with fendiline for 24 or 48 hours. Treatment with 1 µM and 5µM fendiline had no significant effect on eVP40 PM at 24 hours post treatment (Fig EV5B,C); therefore, EGFP-mVP40 PM localization was not assessed further at 24 hours. Since no significant change in EGFP-mVP40 PM localization was observed after 48 hours with either 1 µM or 5 µM fendiline treatment (Fig 5A *bottom panel* and Fig 5B), mVP40 was excluded from further experiments. In agreement with our results thus far, 1 μM fendiline did not significantly inhibit EGFP-eVP40 PM localization after 48 hours of treatment (Fig 5A *top panel* and Fig 5C). However, treatment with 5 µM fendiline for 48 hours led to a modest reduction in EGFP-eVP40 PM localization (∼6% reduction compared to control treated cells; not significant as p=0.08) (Fig 5A *top panel* and Fig 5C). However, the reduction of eVP40 PM localization was not robust enough to lead to the observed inhibition of EBOV by fendiline treatment in our BSL-4 studies (Fig 4). One possible explanation is that a limitation of this technique is the inability to differentiate the extent of VP40 oligomerization occurring using basic confocal microscopy. Therefore, it is possible that fendiline reduced PS levels within the PM, but not significantly enough to block VP40’s ability to bind to the PM.

**Figure 5.**
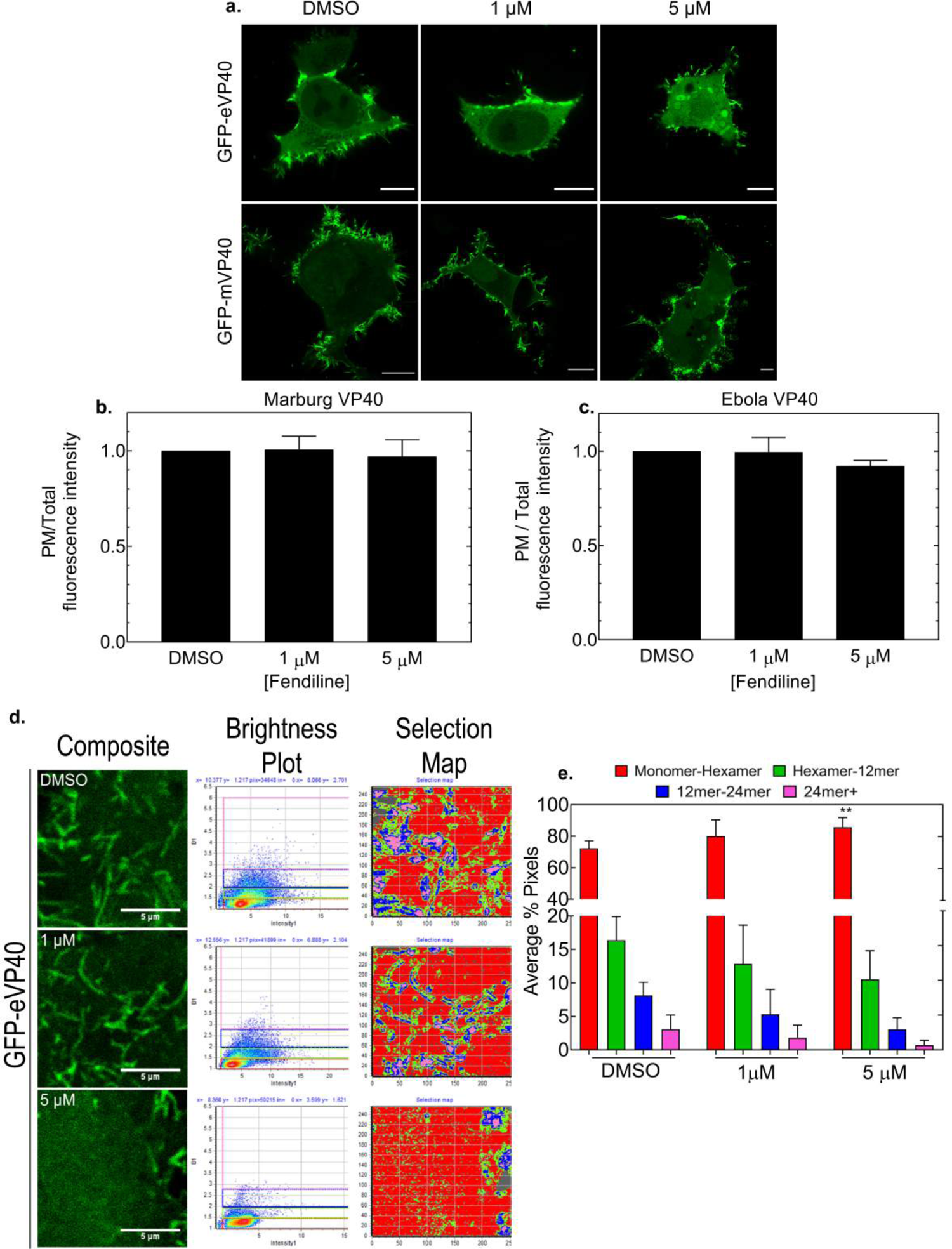
Analysis of eVP40 and mVP40 cellular localization and oligomerization following fendiline treatment. **A-C** Effect of fendiline on eVP40 and mVP40 PM localization in HEK293 cells after 48 hours of treatment. **A** Representative confocal images from live cell imaging experiments of HEK293 cells expressing EGFP-WT-eVP40 (top panel) and EGFP-WT-mVP40 (bottom panel) after 48 hours of fendiline treatment. scale bars= 10 µm. Effect of fendiline on eVP40 **(B)** and mVP40 **(C)** PM localization was quantified by the ratio of EGFP fluorescence intensity at the PM / total EGFP fluorescence intensity (and normalized to DMSO control). N>15, n=3. Values are reported as mean ± s.d. A one-way ANOVA with multiple comparisons was performed compared to the DMSO control. **D-E** Analysis of eVP40 oligomerization in HEK293 cells in response to 48 hour fendiline treatment using N&B analysis. **D** *Left panel:* Representative images from time-lapse (30 frames) of HEK293 expressing EGFP-WT-eVP40 and treated with fendiline for 48 hours. scale bar = 5 µm. *Middle panel:* Brightness and Intensity plots for each representative image. *Right panel:* Selection map correlating each pixel in the representative image to an oligomerization state (b value) (red: monomer-hexamer, green: hexamer-12mer, blue: 12mer-24mer, pink: >24mer). **E** Average % pixel quantification from panel (d)= % of GFP-WT-eVP40 with brightness values corresponding to monomer-hexamer (∼1.-1.6), hexamer-12mer (∼1.6-2.0), 12mer-24mer (2.0-3.2) and >24mer (>3.2) over the total pixels within each image. Values are reported as mean ± s.d.; N≥9, n=3; A two-way ANOVA was performed with Dunnett’s multiple comparisons compared to the control DMSO % average pixels (**p=0.0035). PM: plasma membrane; N&B: Number & Brightness analysis.

### VP40 oligomerization is significantly reduced by fendiline treatment

PS is also a key factor promoting the self-assembly of VP40 into the matrix layer of the budding virion (Adu-Gyamfi *et al*, 2013; Adu-Gyamfi *et al*, 2015). This self-assembly process has been highlighted in our *in vitro* crosslinking data (Fig 2D,E) as well as previously reported in live cells utilizing the N&B technique (Adu-Gyamfi *et al*, 2012). To assess how fendiline impacted eVP40 oligomerization in cells, we examined the oligomerization profile of EGFP-eVP40 using the previously described N&B (Adu-Gyamfi *et al*, 2012; Adu-Gyamfi *et al*, 2015; Johnson *et al*, 2016) (Fig 5D,E). The crystal structure, biochemical analysis, and recent Cryo-ET of eVP40 at the PM strongly suggests eVP40 binds to the PM as a dimer (Bornholdt *et al*, 2013; Del Vecchio *et al*, 2018; Wan *et al*, 2020). eVP40 dimers subsequently oligomerize into larger oligomers via CTD-CTD interactions between dimers, leading to the eVP40 matrix layer formation (Wan *et al*, 2020). Therefore, for our data to coincide with the current models of eVP40 oligomerization, EGFP-eVP40 oligomers were grouped into bins based on multiples of the dimer. Since analyzing eVP40 oligomerization by an increase in every dimer leads to an abundance of data as well as partial overlap in fluorescence signal for each dimer addition, we simplified the data presentation and analysis to different bins of eVP40 oligomers (i.e. dimer-hexamer, hexamer-12mer, 12mer-18mer, and >18mer). The average percentage of pixels in each bin was calculated and plotted for HEK293 cells expressing EGFP-eVP40 and treated with either the control or indicated concentration of fendiline for 48 hrs (Fig 5D,E).

Large eVP40 oligomeric structures corresponding to each bin size were readily detectable at the PM in control treated cells (Fig 5D *top panel* and Fig 5E), with ∼72% of eVP40 found as a dimer-hexamer, ∼16% as a hexamer-12mer, ∼8% as a 12mer-18mer, and 3% in complexes >18mer. Treatment with 1 µM fendiline led to a ∼8% increase in dimeric-hexameric eVP40, and small decreases in the larger oligomeric structures, although no changes were statistically significant (Fig 5D *middle panel* and Fig 5E). However, the oligomeric profile of eVP40 was statistically different when cells were treated with 5 µM fendiline. Following 5 µM fendiline treatment there was a significant increase in eVP40 found in the dimer-hexameric state (∼13% increase; **p=0.0035), which was counterbalanced by an equal reduction in the larger oligomeric states (∼6% reduction for hexamer-12mer, 5% reduction for 12mer-18mer, and ∼3% reduction for eVP40 structures >18mer) (Fig 5D *bottom panel* and Fig 5E). These results support our hypothesis that by reducing PS concentration and therefore the pool of PS available for clustering, eVP40 is unable to properly oligomerize once it traffics and binds to the PM. This, in combination with the modest reduction in eVP40 PM binding following fendiline treatment, may therefore impact the production of viral particles as suggested from our BSL-4 studies.

### Fendiline reduced VLP production at the plasma membrane

As fendiline reduced VP40 oligomerization, we sought to determine the effect of fendiline treatment on VLP production using functional budding assays. No significant effect on VLP production was observed for cells treated with 0.5 µM or 1 µM fendiline at either 24 (Fig 6A *lane 3,4* and Fig 6B) or 48 hours post-treatment (Fig 6C *lanes 3,4* and Fig 6D). However, treatment with one dose of 5 µM fendiline for 24 hours led to a ∼25% reduction in VLP production (Fig 6A *lane 5* and Fig 6B) compared to DMSO treated cells (Fig 6A *lane 2* and Fig 6B). More importantly, this reduction in VLP production was even more robust when monitored at 48 hours post-treatment, with a statistically significant ∼60% reduction in the relative budding efficiency of 5 µM fendiline treated cells (*p-0.0260) (Fig 6C *lane 5* and Fig 6D). The reduction in VLPs is supported by our previous findings that a single dose of 5 µM fendiline reduced PS levels, PS clustering and the extent of eVP40 oligomerization at the PM. Therefore, we hypothesize that reduced virus budding is at least partially responsible for fendiline efficacy in authentic EBOV studies (Fig 4).

**Figure 6.**
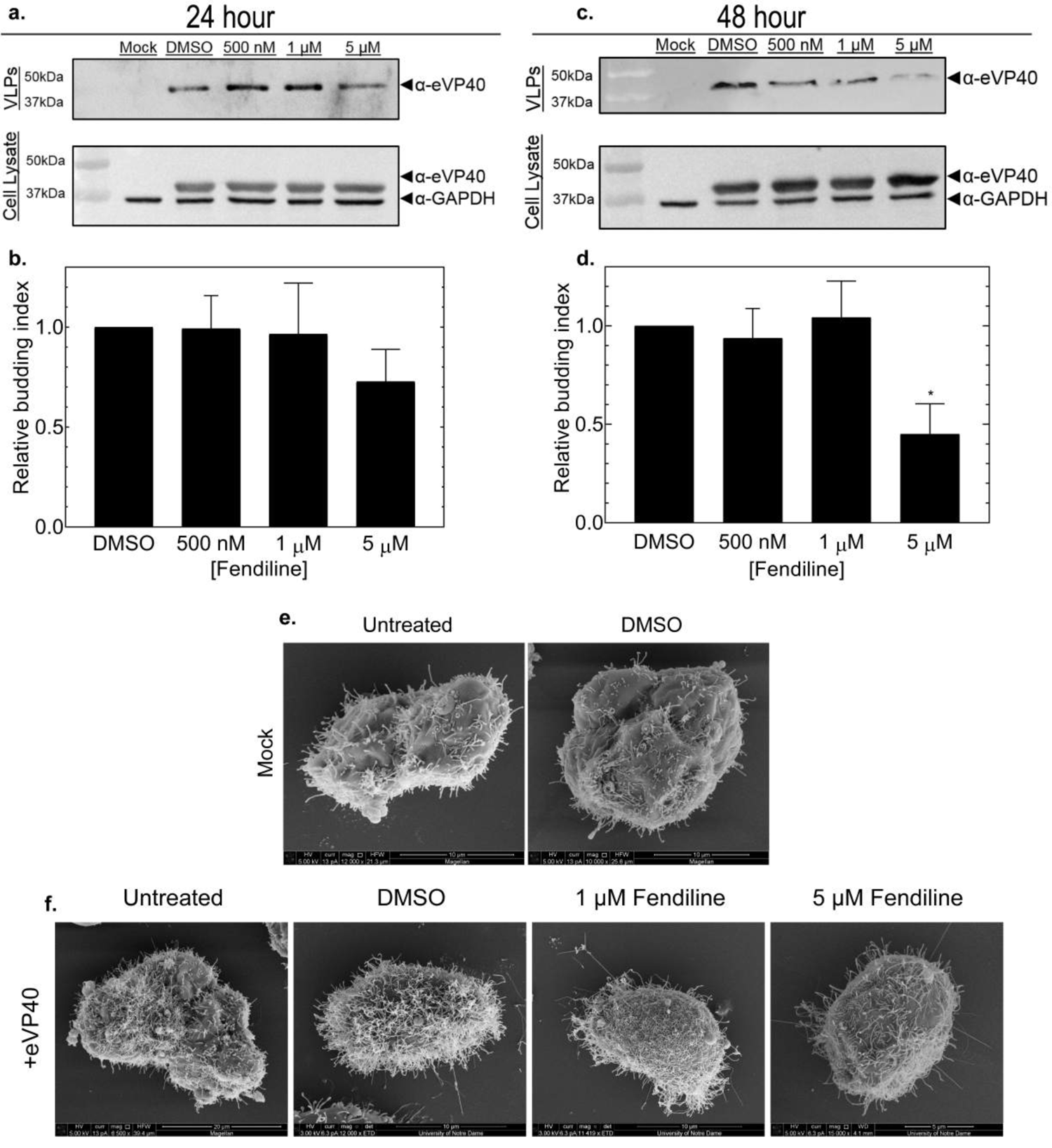
VLP production and morphology in HEK293 cells in the presence of fendiline. **A-D** Functional budding assays assessed at 24 hours (A-B) and 48 hours (C-D) post treatment. **A** Representative western blot of budding assays performed at 24 hours. VLP samples (top panel) and cell lysate samples (bottom panel) collected from HEK293 cells and immunoblotted for eVP40 expression; GAPDH served as a loading control. eVP40 detected by (Rabbit α-eVP40 and HRP-Goat α-Rabbit); GAPDH detected by mouse α-GAPDH and HRP-Sheep α-Mouse) **B** Quantification of relative budding index at 24 hours post fendiline treatment. Relative budding index was determined by the western blot band density of eVP40 in the VLP fraction/(total eVP40 cell lysate + eVP40 VLP band density) and was normalized to the DMSO control. Cell lysate eVP40 band density was normalized to GAPDH band density prior to use in budding index quantification. n=3. Values are reported as mean ± s.d. A one-way ANOVA was performed with multiple comparisons compared to the DMSO control. **C** Representative western blot of budding assays performed at 48 hours. VLP samples (top panel) and cell lysate samples (bottom panel) collected from HEK293 cells and immunoblotted for eVP40 expression; GAPDH served as a loading control. eVP40 detected by (Rabbit α-eVP40 and HRP-Goat α-Rabbit); GAPDH detected by (Mouse α-GAPDH and HRP-Sheep α-Mouse) **D** Quantification of relative budding index at 48 hours post fendiline treatment. Relative budding index was determined by the western blot band density of eVP40 in the VLP fraction/(total eVP40 cell lysate + eVP40 VLP band density) and was normalized to the DMSO control. Cell lysate eVP40 band density was normalized to GAPDH band density prior to use in budding index quantification. n=3. Values are reported as mean ± s.d. A one-way ANOVA was performed with multiple comparisons compared to the DMSO control. (*p=0.0260) **E-F** SEM micrographs of HEK93 cells. **E** Representative micrographs of mock transfected HEK293 cells harvested after 48 hours of no treatment or DMSO treatment. **F** Representative micrographs of HEK293 cells expressing FLAG-eVP40 and harvested after 48 hours of no treatment, DMSO treatment, or the indicated concentration of fendiline. VLPs: virus like particles; SEM: scanning electron microscopy; GAPDH: glyceraldehyde 3-phosphate dehydrogenase; HRP: horseradish peroxidase.

To further investigate the reduction of VLP production in fendiline treated cells, and to determine if there were any observable morphological changes in VLPs, scanning electron microscopy (SEM) experiments were performed (Fig 6E-F). Micrographs of cells expressing FLAG-eVP40 and treated with 5 µM fendiline revealed minimal VLP production at the PM compared to control treated cells (Fig 6F). These findings support the hypothesis that fendiline treatment considerably reduces the production of EBOV VLPs.

### VLP morphology is altered by fendiline treatment

The structure and stability of filoviruses is derived from the VP40 matrix underlying the lipid envelope of virions. Therefore, we utilized transmission electron microscopy (TEM) of purified VLPs to determine if disturbing matrix assembly and altering the lipid components of the PM with fendiline treatment changed VLP morphology and possibly infectivity (Fig 7A-C). During filoviral entry, surface exposed GP and viral envelope PS interact with the receptor T-cell immunoglobulin receptor-1 (TIM-1) (Moller-Tank *et al*, 2013; Brunton *et al*, 2019). To recapitulate entry-competent VLPs (eVLPs), we co-expressed eVP40 with the Ebola virus glycoprotein (eGP). We performed TEM of eVLPs purified from control and 5 µM fendiline treated cells and used ImageJ software to analyze VLP length and diameter (Fig 7A-C). Control eVLPs were heterogenous in length with a mean length of 4.1 µm ± 2.9 (Fig 7A *left panel* and Fig 7B). Control eVLPs diameter also exhibited a level of heterogeneity but had a fairly consistent diameter of 75 nm ± 12.9, which is similar to previous studies of both virions and VLPs (Noda *et al*, 2002; Liu 2014) (Fig 7A *left panel* and Fig 7C). The length and diameter of eVLPs derived from 5 µM fendiline treated cells were significantly less than control eVLPs. Strikingly, fendiline treatment reduced eVLP length by ∼35%, from 4.1 µm to 2.7 µm (*p=0.0139) (Fig 7A *right panel* and Fig. 7B) and modestly reduced eVLP diameter (*p=0.043) (Fig 7A *right panel* and Fig 7C). To ensure that eVLPs derived from fendiline treated cells were not more susceptible to damage during the purification, circular dichroism thermal melting was performed and no difference in eVLP stability was observed (Appendix Fig S1). Reduced eVLP length and diameter could translate into reduced infectivity (e.g., less PS and less surface area and membrane available to bind TIM-1), therefore we next sought to determine the effect of fendiline on eVLP entry.

**Figure 7.**
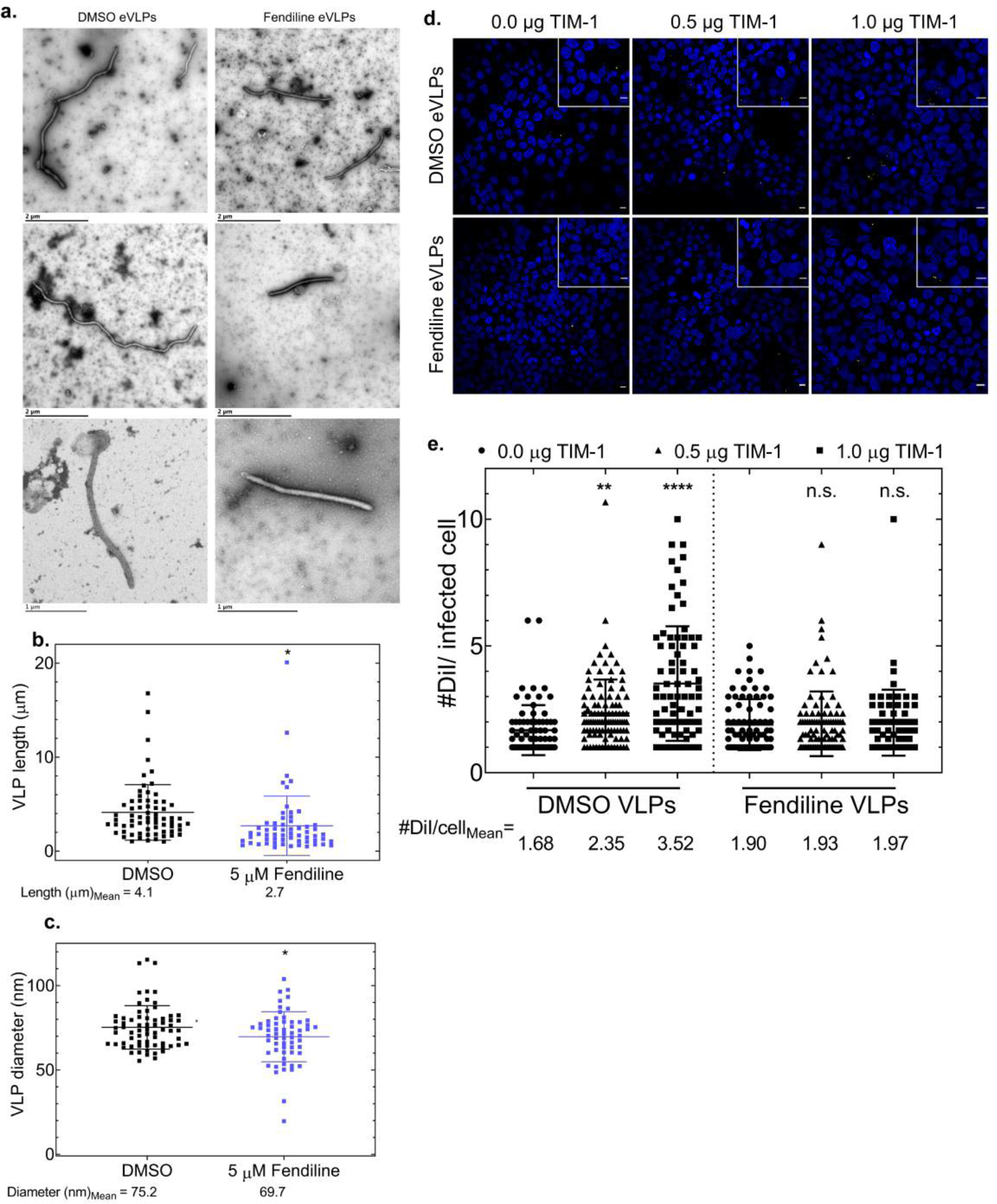
Effect of fendiline on eVLP morphology and TIM-1 dependent eVLP entry. **A-C** TEM analysis of eVLP morphology. **A** Representative transmission electron micrographs of eVLPs purified from HEK293 cells expressing FLAG-eVP40 and eGP following 48 hours of DMSO (left panel) or 5 µM fendiline treatment (right panel). **B** Quantification of eVLP length (µm) of DMSO-derived eVLPs (black) and fendiline-derived eVLPs (blue). N>50, n=3. Values are reported as mean ± s.d. A two-tailed t-test was performed (**p=0.0139). **C** Quantification of eVLP diameter (nm) of DMSO-derived eVLPs (black) and fendiline-derived eVLPs (blue). N>50, n=3. Values are reported as mean ± s.d. A two-tailed t-test was performed (*p=0.0430). **D-E** Fluorescence based DiI TIM-1 dependent entry assay. **D** Representative confocal images from the DiI-entry assay comparing entry of eVLPs produced from DMSO (top panel) and fendiline-treated HEK293 cells (bottom panel) into target cells (HEK293 cells transiently expressing increasing amounts of TIM-1; 0.0 µg, 0.5 µg, 1.0 µg). A stack of 10 frames was acquired for each image. DiI (initially red) was recolored to yellow for easier observation in print; blue (Hoechst 3342 stain); scale bar = 10 µm. **E** Quantification of eVLP entry was performed by calculating the total number of DiI punctate / the total number of DiI-positive cells. Three images from each z-stack was quantified. N=9, n=3. Values are reported as mean ± s.d. A one-way ANOVA was performed with multiple comparisons against the 0.0 µg TIM-1 condition for both DMSO- and fendiline derived eVLPs.(****p<0.0001; **p=0.0093). eVLP: entry-competent viral like particles; TEM: transmission electron microscopy; TIM-1: t-cell immunoglobulin receptor-1; eVLPs: entry-competent VLPs; eGP: Ebola glycoprotein; DiI: 1,1’-Dioctadecyl-3,3,3’,3’-Tetramethylindocarbocyanine Perchlorate.

### Fendiline blocks EBOV eVLP entry

A common characteristic of viral infectivity is the relationship between virion associated PS and the TIM-1 receptor on target cells (Kondratowicz *et al*, 2011; Moller-Tank *et al*, 2013; Wang *et al*, 2017; Dejarnac *et al*, 2018; Brunton *et al*, 2019). Moreover, it has been previously reported that other ASM inhibitors blocked EBOV infectivity (Miller *et al*, 2012). To determine if fendiline treatment reduced the entry of eVLPs, we performed a fluorescent based entry assay using 1,1’-dioctadecyl-3,3,3’,3’-tetramethylindocarbocyanine perchlorate (DiI) labelled eVLPs (Nanbo *et al*, 2010; Kuroda *et al*, 2015; Nanbo *et al*, 2019). By testing entry of eVLPs derived from fendiline treated cells rather than the entry of eVLPs on fendiline treated cells, we were able to determine how fendiline treatment affected eVLP entry rather than how inhibition of ASM in target cells affected eVLP entry (as previously described (Miller *et al*, 2012). If entry of the eVLPs was not altered by fendiline treatment, one would expect a dose-dependent increase in infectivity with increasing TIM-1 expression. Conversely, if eVLP entry was inhibited by fendiline treatment, a dose-dependent increase in eVLP entry would not be observed with increasing TIM-1 expression.

TIM-1 overexpression was increased in target cells and a detectable and significant dose-dependent increase in control eVLP entry was observed, by more than 200% in the highest TIM-1 overexpressing cells (compared to no TIM-1 overexpression; Fig 7D *top panel* and Fig 7E). Remarkably, no measurable increase in eVLP entry was observed for fendiline derived-VLPs across any of the TIM-1 overexpressing target cell conditions (Fig 7D *bottom panel* and 7E). From this comparison, these results suggest that the impaired entry of fendiline eVLPs is a result of reduced PS in the viral envelope, either from smaller VLPs or a lower % of PS content. These findings in combination with the observed reduction in VLP formation further substantiate the significant reduction of EBOV infection observed in our live virus studies following fendiline treatment.

### Mathematical model of in vitro experiments

We next used a mathematical model to predict how the effects of fendiline on both viral budding and entry combine to produce the observed effects in the BSL-4 assays. We calibrated our mathematical model (equations 1-3) to experimental data from the budding, entry and cellular infection assays using approaches and parameter settings outlined in the methods and Table 1. Results from our two-phase calibration procedure are shown in Fig 8A-F and Appendix Fig S2. The model captures key features of the data including a progressive increase in percentage of infected cells over time, differences between MOI as well as limited cell death in the first 48 hours of the experiment (Appendix Fig S2).

**Figure 8.**
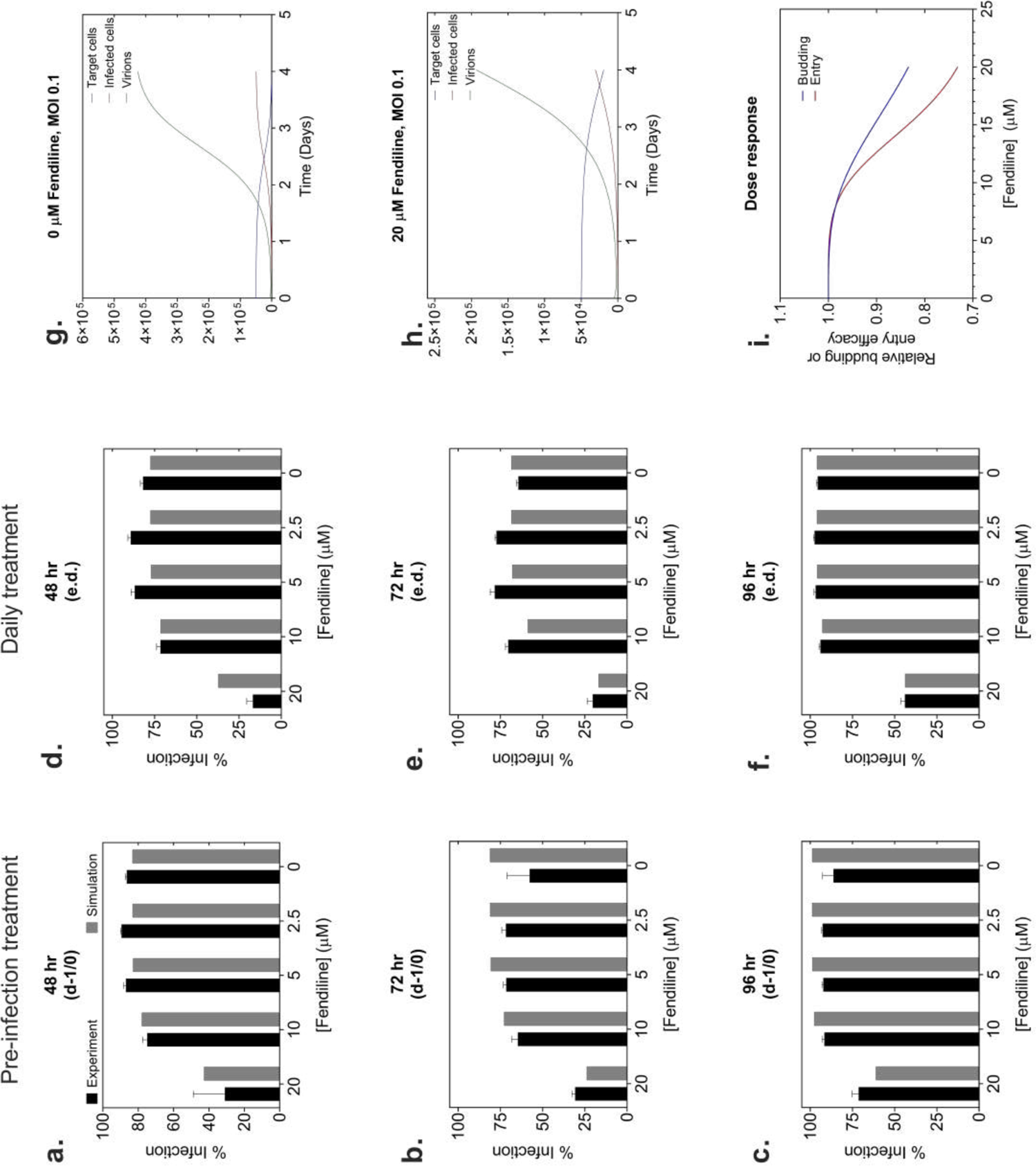
Calibrated mathematical model reproduces key observations in multiple experimental datasets. Percentage infected cells is shown for various fendiline concentrations given prior to infection (d-1/0, **A-C**) or daily (e.d., **D-F**). (**A,D**) MOI 1; (**B,C,E,F**) MOI 0.1. (**G-H**) Model predicted cell and viral dynamics for MOI 0.1. (**i**) Model predicted dose response curves for fendiline effects on viral budding and entry in the BSL4 experiments.

**Table 1.** Mathematical model parameter descriptions and values

| Symbol | Description | Unit | Value from calibration to budding and entry assays | Ranges based on fits to budding and entry assays | Value from calibration to BSL4 data – using ranges defined by budding and entry assays |
| --- | --- | --- | --- | --- | --- |
| $\beta$ | Infection rate constant of target cells by free virions | (Virion.day)<sup>-1</sup> | N/A | 0 - inf | 6.6x10<sup>-6</sup> |
| $\delta$ | Death rate constant of infected cells | Day<sup>-1</sup> | N/A | 0 - inf | 4.6x10<sup>-6</sup> |
| c | Decay rate constant of free virions | Day<sup>-1</sup> | N/A | 0 - 5 | 5 |
| p | Viral production rate constant by infected cells | Virions.(Infected cell.day)<sup>-1</sup> | N/A | 1 - inf | 49 |
| $H_b$ | Hill constant for Fendiline effects on VLP budding | | 2.7 | 0.5 - 7 | 3.4 |
| $H_e$ | Hill constant for Fendiline effects on VLP entry | | 11 | 0.5 - 20 | 4.7 |
| $C_{50}^b$ | Concentration with 50% of maximum Fendiline effect on VLP budding | $\mu\text{M}$ | 5.7 | 2 - 20 | 18 |
| $C_{50}^e$ | Concentration with 50% of maximum Fendiline effect on VLP entry | $\mu\text{M}$ | 4.8 | 0.5 - 20 | 15 |
| $E_{max}^b$ | Maximum Fendiline effect on VLP budding | | 0.98 | 0.3 - 1 | 0.3 |
| $E_{max}^e$ | Maximum Fendiline effect on VLP entry | | 0.76 | 0.3 - 1 | 0.33 |

The dynamics behind these calibrated figures suggested that fendiline treatment significantly delayed the infection process (Fig 8G,H), resulting in the observed decrease in percent infected cells with treatment over 4 days. The effects of fendiline on budding and entry are estimated to have similar pharmacodynamics (PD), with entry effects estimated to have a slightly stronger response (lower C_50_ and higher E_max_) compared to budding (Table 1). Based on PD parameters, the response to fendiline was estimated to be weaker in the BSL-4 assays, as is evident by higher C_50_ values and lower E_max_ values compared to the budding and entry assays (Table 1, Fig 8I). These PD parameter differences between budding and entry assays vs BSL-4 results could suggest that other parts of the viral life cycle not affected by fendiline (not quantified explicitly in these experiments) become rate limiting in the BSL-4 assays, thereby reducing the overall effect of fendiline on infection progression. In summary, a mathematical model consistent with three independent experimental systems, predicts a combination of budding and entry effects resulting in the observed BSL-4 effects, and estimates PD parameters for each mechanism.

## Discussion

The host cell PM is exploited by filoviruses for their assembly and budding, where they can egress the host cell to form a new virion. The matrix protein, VP40, is the main driver of this process as it harbors a high affinity for lipids in the PM inner leaflet. Lipid binding by VP40, which includes selectivity for PS (Ruigrok *et al*, 2000; Scianimanico *et al*, 2000; Adu-Gyamfi *et al*, 2015; Del Vecchio *et al*, 2018) and PI(4,5)P_2_ (Johnson *et al*, 2016; Johnson *et al*, 2018) drives and stabilizes, respectively, VP40 oligomers that are necessary for viral budding. In fact, VP40 has been shown to be sufficient (in the absence of other filovirus proteins) to form VLPs from the host cell PM that are nearly indistinguishable from virions (Jasenosky *et al*, 2001; Noda *et al*, 2002; Licata *et al*, 2004). Again, this underscores the unique properties of VP40 structure and sequence, which provides a template for host lipid binding, oligomerization, and sufficient information to encode cues for scission to complete the viral budding process. Further, VP40 derived VLPs enter cells in a PS-dependent manner despite the absence of the EBOV glycoprotein (Jemielity *et al*, 2013). While some of the basic principles between VP40 and PM interactions have been previously revealed, the mechanistic consequences of the interactions and their potential pharmacological targeting have remained unknown.

In this study, we demonstrated that VP40 was able to cluster PS *in vitro* and in the PM of cells. VP40-dependent PS clustering required PS binding and efficient PM localization of VP40, as the K224A-VP40 mutant, previously defined as a PS-binding residue did not significantly increase PS clustering. VP40 oligomerization was also crucial to PS clustering efficiency as a VP40 oligomerization deficient mutant, which can still exhibit PM localization, significantly reduced PS clustering. This is an important point as the PM of host cells is generally thought to harbor 20-30 mol% PS in the inner leaflet. In fact, VP40 effectively binds and oligomerizes on PS-containing membranes with compositions close to the PS-content of the PM inner leaflet. In contrast, PS concentrations below 15 mol% didn’t provide robust affinity and oligomerization compared to those with 22 mol% PS and greater. PS clustering and PS content may also play a critical role in the loss of asymmetry that occurs during the EBOV budding process. For instance, pooling of PS in distinct regions of VP40 assembly may provide a cue for scramblases shown to distribute PS to the outer leaflet of the PM during the viral budding process (Nanbo *et al*, 2010; Younan *et al*, 2018).

A recent study demonstrated that silencing PSS1, an enzyme responsible for PS synthesis in mammalian cells was sufficient to inhibit EBOV replication (Younan *et al*, 2018). However, to the best of our knowledge, small molecules aimed at targeting host cell lipid distribution have not previously been tested against EBOV. Fendiline was a logical choice to form an initial hypothesis of an FDA-approved drug that could inhibit EBOV budding as it was recently shown to lower PM PS (Cho *et al*, 2016) and inhibit K-Ras signaling (van der Hoeven *et al*, 2013; van der Hoeven *et al*, 2018) suggesting this FDA-approved drug may be sufficient to inhibit EBOV budding. Indeed, in BSL-4 experiments, fendiline was able to inhibit EBOV replication >75% at 20 µM when given every day post-infection. Follow up mechanistic studies demonstrated that fendiline efficacy was due to inhibition of VP40 oligomerization, viral budding and viral entry. Thus, fendiline, which reduced the PS content of HEK293 cells by ∼30%, subsequently reduced PS clustering and VP40 oligomerization necessary for efficient viral budding. VLPs that did form from fendiline-treated cells had an overall reduced length and surface area, which likely combined with the reduced PS-content of the virus or VLPs to limit subsequent viral entry. Thus, disruption of PM PS content by one small molecule was sufficient to effect at least three important steps in the filovirus life cycle. Importantly, an increase in intracellular calcium due to fendiline can be ruled out in reducing and/or inhibiting EBOV and VP40 budding as calcium has been shown to be a critical component of VLP formation and virus replication (Han Z *et al*, 2015). Thus, an increase of calcium in HEK293 cells would be expected to increase and not decrease budding as we’ve observed.

Overall, this study lends credence to the hypothesis that host processes may be targeted to inhibit viral replication and spread. While the potency of fendiline may be low, the combination of fendiline with other FDA-approved drugs that have shown efficacy against EBOV (Johansen *et al*, 2013; Johansen *et al*, 2015; Nelson *et al*, 2016; Dyall *et al*, 2018; Jasenosky *et al*, 2019) hold further promise. Fendiline was effective at 5 µM for reducing PS content in cells and the relationship to fendiline concentrations used *in vivo* is an important point to address in the future. Fendiline has been used in both single (200-1200 mg) and multiple doses (400 mg, twice) per day, which was followed with pharmacokinetic evaluation yielding maximal plasma levels in patients of 9-170 ng/mL (Weyhenmeyer *et al*, 1987). At the upper range of fendiline detected in the plasma, 170 ng/mL = 540 nM fendiline. Thus, at the upper value of fendiline detected from oral dosing, the concentration was lower than the 5 µM used to lower cellular PS. Part of the issue in fendiline dosing is rapid metabolism with a bioavailability of ∼20% depending on the mg dose given (Kurovetz *et al*, 1982). Thus, the issue isn’t necessarily fendiline toxicity and efficacy but drug metabolism of fendiline in the plasma and through the hepatic 1^st^ pass effect that limit plasma concentration available (Kurovetz *et al*, 1982). Thus, future studies geared towards more stable drugs that can alter PS content through a similar mechanism of action (via ASM inhibition) may be beneficial.

Recently, fendiline analogs have been generated that have nanomolar potencies against K-Ras PS-dependent plasma membrane localization and were effective at inhibiting cancer cell proliferation in the low micromolar range (Wang *et al*, 2021). Fendiline has also been shown to block SARS-CoV-2 S-mediated fusion (Xiao *et al*, 2020) and cause *Leishmania infantum* death via mitochondrial depolarization (Reimão *et al*, 2016). Thus, our work and these studies lay a framework to improved pharmacological targeting strategies against either VP40 matrix assembly or PS clustering that would not only reduce viral budding and spread, but lower subsequent viral entry, which partially relies on PS in the viral envelope (Moller-Tank *et al*, 2013; Jemielity *et al*, 2013; Moller-Tank *et al*, 2014; Brunton *et al*, 2019). Notwithstanding pharmacological principles learned from this study, a critical balance between VP40 and PS has been resolved demonstrating a critical need for VP40 clustering in the assembly and budding process. VP40 oligomers are needed for enhanced PS clustering where PS clustering seems to ensure optimal VP40 oligomerization.

## Material and Methods

### Reagents & solutions

PBS, DMEM, ionomycin and Lipofectamine^TM^ LTX with Plus^TM^ Reagent, and Invitrogen^TM^ Molecular Probes^TM^ Hoechst 3342 were purchased from Fisher Scientific, heat-inactivated fetal bovine serum (FBS) was purchased from Hyclone, and Minimum Essential Medium (MEM) was purchased from Corning. Invitrogen Live Cell Imaging Solution, DiI Stain, Halt^TM^ Protease Inhibitor Cocktail, Pierce BCA Assay kit, and BS3 were purchased from ThermoFisher Scientific. Non-essential amino acids (NEAA) were purchased from Sigma Aldrich and L-glutamine was purchased from Gibco. Alexa Fluor^TM^ 488 C_5_ Maleimide (Alexa-488) for protein conjugation was purchased from Invitrogen. Fendiline was purchased from Cayman Chemical Company, prepared in DMSO and stored at −20 °C. Ultra-Pure Grade DMSO was purchased from VWR and the Ni-NTA slurry was purchased from Qiagen. L1 chips for SPR experiments were purchased from GE Healthcare. For cell viability assays, CellTiter-Glo® was purchased from Promega. Antibody information for immunoblotting and immunofluorescence can be found in Table 2. Ten percent neutral buffered formalin was purchased from Val Tech Diagnostics (Brackenridge, PA). Cell staining buffer was purchased from BioLegend. stain was purchased from Fisher Scientific.

### Plasmids

EGFP, EGFP-eVP40 and EGFP-WE/A-eVP40 were prepared as described previously (Adu-Gyamfi *et al*, 2012; Johnson *et al*, 2016). GFP-K224A-eVP40 was prepared by site directed mutagenesis (Johnson *et al*, 2018) GFP-mVP40 was used as described previously (Wijesinghe & Stahelin 2015; Amiar *et al*, 2021). The GFP-LactC2 plasmid was a kind gift from Sergio Grinstein (University of Toronto). GFP-PLCδPH was a kind gift from Tamas Balla (NIH). pCAG-GPI-GFP was a gift from Anna Katerina Hadjantonakis (Addgene #32601). pEGFP-N3-Annexin A2 was a gift from Volker Gerke & Ursula Rescher (Addgene #10796). pCAGGS-FLAG-eVP40 (NR49337) and pcDNA3.1-eGP (NR-19814) were obtained from BEI Services. pCAGGS-TIM-1 was from Heinz Feldmann (Watt *et al*, 2014).

### Lipids and LUV preparation

POPC (#850457), DPPC (#850355), cholesterol (#700000), DPPS (#840037), POPE (#850757), POPS (#840034), Brain PI(4,5)P_2_ (#840046), and TopFluor® TMR-PS (#810242) were purchased from Avanti Polar Lipids, Inc. (Alabaster, AL) and stored in chloroform and/or methanol at −20°C until use.. For large unilamellar vesicle (LUV) preparation used in SPR and chemical crosslinking experiments, lipid mixtures were prepared at the indicated compositions (using a POPC matrix as the background and control and the indicated additions of PS at the expense of POPC), dried down to lipid films under a continuous stream of N_2_, and stored at −20°C until further use. On each day of experiments, LUVs were brought to room temperature, hydrated in either SPR buffer (10 mM HEPES, 150 mM NaCl, pH 7.4) or chemical crosslinking buffer (260 µM Raffinose pentahydrate in PBS, pH 7.4), vortexed vigorously, and extruded through a 100 nm (SPR experiments) or 200 nm (chemical crosslinking experiments) filter. Vesicle size was confirmed by dynamic light scattering using a DelsaNano S Particle Analyzer (Beckman Coulter, Brea, CA).

### Preparation and imaging of GUVs

GUVs were prepared by a gentle hydration method (Reeves *et al*, 1969; Darszon *et al*, 1980; Yamashita *et al*, 2002). Briefly, 1 mM of lipid control mixture was made and contained 1,2-dipalmitoyl-sn-glycero-3-phosphocholine (DPPC), cholesterol (Chol) and fluorescent phosphatidylserine (TopFluor® TMR-PS) at 90:9.8:0.2% molar ratio. For PS clustering and eVP40 binding analysis, 1,2-dipalmitoyl-sn-glycero-3-phosphoserine (DPPS) alone or with brain phosphatidylinositol 4,5-bisphosphate PI(4,5)P_2_ were added at 40 and 2.5% molar ratios, respectively, and the ratios of DPPC were adjusted accordingly. The lipid mixtures were prepared in 5 mL round-bottom glass flasks and the chloroform was removed with rotary movements under a continuous stream of N_2_ and even heat at 55°C until complete evaporation of the solvent. The lipid films were then hydrated overnight at 50°C (DPPC:Chol:TopFluor TMR-PS) and 55°C (DPPC:Chol:DPPS:PI(4.5)P_2_:TopFluor TMR-PS) in an appropriate volume of GUV hydration buffer (150 mM NaCl, 10 mM HEPES, 0.5 M sucrose, pH 7.4).

For imaging, the freshly hydrated GUVs were diluted 10 times in GUV dilution buffer (150 mM NaCl, 10 mM HEPES, 0.5 M glucose, pH 7.4) and placed on a 6-mm diameter chamber made from a silicon sheet using a core sampling tool (EMS # 69039-60). The silicon chamber was mounted on a 1.5-mm clean coverglass (EMS # 72200-31) pre-coated with 1 mg/mL BSA. The set up was then assembled in an Attofluor chamber (Invitrogen # A7816) and 1.25 µM eVP40-Alexa488 was added. GUVs imaged for eVP40 and PS colocalization and clustering analysis was performed at 37°C on a Nikon Eclipse Ti Confocal inverted microscope (Nikon Instruments, Japan), using a Plan Apochromat 60x 1.4 numerical aperture oil objective and a 100x 1.45 numerical aperture oil objective, respectively. A 488 nm argon laser was used to excite GFP and a 561 nm argon laser was used to excite TopFluor®-TMR-PS. The 3D reconstruction was performed using ImageJ. Mander’s correlational analysis was performed using the plugin JACoP (Bolte & Cordelières 2006).

### Cell culture, transfections, pharmacological treatments

All BSL-2 studies were performed using HEK293 cells obtained from the American Type Culture Collection and cultured in DMEM supplemented with 10% FBS and 1% PS. Transient transfections were performed using Lipofectamine^TM^ LTX + PLUS^TM^ Reagent, according to the manufacturer’s protocol. All transfections were performed in DMEM supplemented with 10% FBS. Treatment with fendiline (in DMSO) occurred at 5-hours post-transfection in DMEM supplemented with 10% FBS. BSL-4 assays were also performed using Vero E6 cells cultured in MEM, 5% heat-inactivated fetal bovine serum, 1% L-glutamine, and 1% NEAA. Both HEK293 and Vero E6 cells were cultured and incubated at 37°C, 5% CO_2_, 80% humidity.

### Immunoblotting

Samples prepared for western blotting analysis were first separated using SDS-PAGE (8% for chemical crosslinking and 12% for cell lysates and VLPs). Following transfer onto a nitrocellulose membrane, membranes were blocked with 5% MILK-TBST and analyzed with their respective antibodies (See Table 2). Antibodies were detected using an ECL detection reagent and imaged on the ImageQuant LAS 4000 or Amersham Imager 600 (GE Healthcare Life Sciences). All quantitative analysis derived from western blotting was performed using densitometry analysis in ImageJ.

**Table 2.** Antibodies used in immunoblotting

| Target | Species | Tag | Dilution | Cat. # | Source |
| --- | --- | --- | --- | --- | --- |
| eVP40 | Rabbit |  | 1/40,000 | 0301-010 | IBT Bioservices |
| Rabbit | Goat | HRP | 1/5,000 | 205718 | Abcam |
| GAPDH | Mouse |  | 1/5,000 | 8245 | Abcam |
| Mouse | Sheep | HRP | 1/10,000 | 6808 | Abcam |
| EBOV-GP (KZ52) | Human | | 1.0 $\mu\text{g/ml}$ | 0260-001 | IBT Bioservices |
| Human | Goat | Alexa-488 | 0.7 $\mu\text{g/ml}$ | A11013 | Invitrogen |
| Mouse | Goat | Alexa-488 | 1.0 $\mu\text{g/ml}$ | A11029 | Invitrogen |
| 6xHis | Mouse |  | 1/1,000 | A5588 | Sigma |

### Protein purification and Alexa-488 C_5_ Maleimide labelling

The His_6_-eVP40-pET46 expression vector was a kind gift from Erica Ollmann Saphire (La Jolla Institute for Immunology) and was expressed and grown in Rosetta^TM^ 2 BL21 DE3 competent cells (Merck Millipore, Billerica MA). The pet28a-His_6_-Lact C2 bacterial expression plasmid was a kind gift from Dr. Sergio Grinstein. His_6_-eVP40 and His_6_-LactC2 were grown and purified as described previously (Johnson *et al*, 2016). Following elution from a Ni-NTA slurry (Qiagen), the protein samples were then further purified using size exclusion chromatography on a HiLoad 16/600 Superdex 200 pg column (ÄKTA pure, GE Healthcare). The desired fractions containing dimeric VP40 or monomeric LactC2 were collected, concentrated and stored in 10 mM Tris, 300 mM NaCl, pH 8.0. Protein concentration was calculated using the Pierce BCA assay and the protein was stored at 4°C for no longer than 14 days.

Labeling of eVP40 cysteine residues was carried out using Alexa-488-C_5_-maleimide. eVP40 dimer was treated with a 1.5-fold molar excess of Alexa-488-C5-maleimide dissolved in DMSO for 2 hours at room temperature in maleimide labelling buffer (20 mM NaPi solution pH 7.4, 150 mM NaCl, 4 M Guanidine HCl). The labeling reaction was quenched by adding diothiothreitol (final concentration of 50 mM) and the labeled protein was separated from the non-conjugated dye on a HiLoad 16/600 Superdex 200 pg column. Fractions containing eVP40 dimer were collected and concentrated. The labelling efficiency and protein concentration were estimated using a NanoDrop according to Invitrogen’s instructions.

### Plasma membrane localization confocal microscopy

Live cell imaging experiments were performed at 24 hours and 48 hours post treatment. Experiments to quantify fluorescent protein PM localization were performed on a Zeiss LSM 710 inverted microscope using a Plan Apochromat 63x 1.4 numerical aperture oil objective. A 488 nm argon laser was used to excite GFP/EGFP. GFP-LactC2 PM localization was quantified ratiometrically by comparing the PM signal vs. the cytosolic signal. GFP-EBOV-VP40 PM localization was quantified ratiometrically by comparing the PM signal vs. the total fluorescence signal within the cell.

### TMR-PS quenching experiment

The Top Fluor TMR quenching analysis was performed as described previously (Zhao *et al*, 2010). Briefly, LUVs with 0.5% TopFluor® TMR-PS and increasing amounts of DPPS (0, 1, 5, 10, 20, 40, 60%) with or without PI(4.5)P_2_ (2.5% with 60% DPPS) were made as described above. LUVs were then mixed at a final concentration of 40 µM with corresponding concentrations of eVP40 dimer in 10 mM HEPES, 160 mM NaCl pH 7.4 in a black/clear bottom 96-well plate and incubated at 37°C for 30 min. The Top Fluor TMR was excited at 547 nm and the fluorescence was recorded from 555 to 600 nm with no cutoff wavelength using the plate reader SpectraMax M5e (Molecular Devices, St Jose, CA). The fluorescence quenching ratio, DF, was calculated according to the ratio (F_0_/F), where F_0_ is the fluorescence in absence of protein and F is the fluorescence at a given eVP40 concentration.

### Cellular Top Fluor TMR-PS clustering confocal microscopy

Each experimental day, a 100 μM working stock of TopFluor® TMR-PS in methanol was prepared. Immediately prior to imaging, cells were placed in 4°C for 5 min. The working stock was diluted to a final 500 nM TopFluor® TMR-PS solution in 3 mg/mL BSA/PBS. The 500 nM TopFluor® TMR-PS/BSA/PBS solution was incubated with cells at 4°C for 10 min, rinsed three times with cold PBS, and immediately imaged in fresh cold PBS. Top Fluor TMR was excited at 560 nm and GFP was excited at 488 nm. For PS clustering analysis, a custom macro in ImageJ was used. Prior to the macro analysis, background was subtracted and the contrast was enhanced. To isolate the PM area, a default threshold was applied. To isolate PS clusters the Moments analysis thresholding (Tsai 1985) was applied. Following the moments analysis thresholding, the custom ImageJ macro was applied: despeckle, close-, fill holes, and remove outliers (radius=5, threshold=50). The sum of the remaining particles area was calculated, as well as the PM area. %PS clustering was calculated according to the ratio (Area_clusters_/Area_plasma membrane_).

### Number & Brightness

Number & Brightness experiments were performed as described previously (Adu-Gyamfi *et al*, 2017; Bobone *et al*, 2017; Johnson *et al*, 2018) on a Zeiss LSM 880 upright microscope using a LD “C-Apochromat” 40x/1.1 W Corr M27 objective. HEK293 cells expressing either GFP, GFP-LactC2 or GFP-eVP40 were treated with fendiline (1 or 5 μM) for 48-hours prior to N&B analysis. Cells were imaged in phenol-free live cell imaging solution. For each experimental day, the brightness value of a monomer was determined in cells expressing monomeric GFP. Each image was acquired using the same laser power (0.01), resolution (256×256), pixel dwell time (16 us), frames (50), and zoom (pixel size of 50 nm). SimFCS Globals Software (Laboratory for Fluorescence Dynamics, University of California, Irvine, CA) was used for analysis.

### Chemical crosslinking

His_6_-eVP40 and His_6_-LactC2 were purified as previously described in *Protein Purification*. LUVs containing POPC and Brain PI(4,5)P_2_ (2.5%) with varying PS mol% composition (0, 15,30, 60%) were prepared as previously described in *Lipids & Vesicle Preparation*. Experimental protocol was adapted from (Johnson *et al*, 2016), and the manufacturers protocol for BS^3^ (ThermoFisher) with each step performed at RT. In brief, protein (final concentration of 0.3 µM in PBS pH 7.4) was mixed with LUVs (final concentration = 660 µM) at a 1:1 volumetric ratio for 30 min. Protein bound LUVs were separated from unbound protein through centrifugation (75,000 x *g*, 30 min, 22°C), and resuspended in PBS (pH 7.4) buffer containing BS^3^ (final concentration of 200 µM). Samples were incubated for 45 min, quenched with glycine for 15 min and then analyzed through western blotting. Following immunoblotting, oligomerization of VP40 was quantified ratiometrically by comparing VP40_o_ vs. VP40_m+d_ (where VP40_o_ is the oligomeric VP40 band density (>75 kDa) and VP40_m+d_ is the sum of monomeric and dimeric eVP40 (∼37 and 74 kDa) band density).

### Surface plasmon resonance

To determine the affinity of His_6_-eVP40 to LUVs with increasing PS concentrations, SPR was performed. SPR experiments were performed at 25°C using a Biacore X100 as described previously (Adu-Gyamfi *et al*, 2015). In brief, an L1 chip was coated at 5 µL/min with LUVs containing 0% PS (100% POPC, PS added at the expense of POPC) on flow cell 1 and either 1%, 11% or 22 mol% POPS on flow cell 2 (LUV preparation described in previous section, *Lipids and Vesicle Preparation)*. The LUV conjugated chip was stabilized by washing with 50 mM NaOH and blocked with 0.1 mg/mL BSA (in SPR buffer) at a flow rate of 10 μL/min until the response on each flow cell was <100 response units (RU). For quantitative affinity analysis, each concentration of eVP40 was injected for 540 s at a flow rate of 10 μL/min with a 180 s delay, and the difference in response between flow cell 1 and flow cell 2 was recorded (ΔRU). The apparent *K*_d_ of vesicle binding was determined using the non-linear least squares analysis: 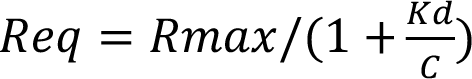 where *R*_eq_ (measured in RU) is plotted against protein concentration (C). *R*_max_ is the theoretical maximum RU response and *K*_d_ is the apparent membrane affinity. Data were fit using the Kaleidagraph fit parameter of (m0*m1)/(m0+m2);m1=1100;m2=1. ΔRU data was normalized in GraphPad Prism 8 for windows (La Jolla, CA) and plotted in Kaleidagraph (Reading, PA).

### Lipidomics

HEK293 cells were treated with the indicated concentration of fendiline for 48 hours, collected through centrifugation, rinsed with PBS and protein concentration was determined. Cells were pelleted, flash frozen in liquid N_2_ and stored at −80°C until subsequent LC/MS/MS processing by Avanti Polar Lipids, Inc.. Prior to LC-MS/MS analysis, lipids were extracted using the Folch method (Folch *et al*, 1957). The bottom chloroform layer taken after centrifugation was diluted with internal standards for LPA, LPS, PA and PS for quantization by injection on LC-MS/MS. Samples were injected on a LC-MS/MS method using a Waters Acquity UPLC / AB Sciex 5500 MS system performing reversed phase separation of LPA and LPS and PA and PS components with MS/MS detection. Each molecular species identified by the [M-H] m/z of its acyl carbon:double bond (CC:DB i.e. 34:2 PA) was quantified against the response of the internal standards of known concentration. Content of individual and total LPA/PA and LPS/PS was reported. Values were corrected to 1×10^6^ cells for all samples.

### BSL-4 immunofluorescence assay

Vero E6 cells were seeded at 2e4 cells/well in black 96-well poly-D-lysine treated plates (Greiner Bio-One Cellcoat®). Twenty-four hours prior to infection, fendiline was diluted in 0.5% DMSO and Vero E6 cell culture media at indicated concentrations and added to cells. An equivalent percentage of DMSO in culture media served as the vehicle control. Following pretreatment, compound was removed, and cells were incubated with Ebola virus (Kikwit) at a multiplicity of infection (MOI) of 0.1 or 1.0 in a BSL-4 located at USAMRIID. Following absorption for 1 hour, virus inoculum was removed and cells were washed. Plates were divided into three post-infection treatment groups (day 0, every day-e.d., every other day-e.o.d.), and received either culture media or freshly prepared fendiline or vehicle control. Cells were then treated daily with freshly prepared compound or left to incubate based on their designated treatment group. At 48 hours (MOI=1.0), 72 or 96 hours (MOI=0.1) post infection, cells were washed with PBS and submerged in 10% neutral buffered formalin for 24 hours prior to removal from the BSL-4 laboratory. Formalin was removed and cells were washed with PBS. Cells were blocked with 3% BSA/PBS cell staining buffer (BioLegend) and incubated at 37°C for 2 hours. Ebola virus GP-specific mAb KZ52, diluted in 3% BSA/PBS, was added to appropriate wells containing infected cells and incubated at room temperature for 2 hours. Cells were washed three times with PBS prior to addition of goat anti-human or goat anti-mouse IgG-Alexa-488 secondary antibody. Following 1-hour incubation with secondary antibody, cells were washed 3 times prior to counterstaining with Hoechst’s stain diluted in PBS. Cells were imaged and percent of virus infected cells calculated using the Operetta High Content Imaging System and Harmony® High Content Imaging and Analysis Software (PerkinElmer).

### VLP collections & functional budding assays

HEK293 cells were transfected and treated with fendiline as described in the previous section, *Cell Culture, Transfection & Pharmacological Treatments*. Budding assays were performed as described previously (Johnson *et al*, 2016; Harty 2018). In brief, VLP containing supernatants were harvested from cells and clarified through low speed centrifugation. Clarified VLPs were loaded onto a 20% sucrose cushion in STE buffer (10 mM TRIS, 100 mM NaCl, 1 mM EDTA, pH 7.6), isolated through ultracentrifugation, and resuspended in either 150 mM ammonium bicarbonate (functional budding assays), 2.5% glutaraldehyde in 0.1 M cacodylate buffer (TEM experiments), STE buffer (entry assays) or 0.1 M phosphate buffer for CD and thermal melting (PB; 0.02 M sodium phosphate monobasic, 0.08 M sodium phosphate dibasic, pH 7.4). VLP samples were stored at −80°C for functional budding assays, −20°C for entry assays or 4°C for TEM and CD analysis.

For functional budding assays, cell lysate samples were harvested and lysed on ice with RIPA buffer (150mM NaCl, 5mM EDTA pH=8, 50mM Tris pH 7.4, 1% Triton-X, 0.1% SDS, 0.5% deoxycholic acid) supplemented with Halt^TM^ Protease Inhibitor Cocktail. Prior to separation on a 12% SDS-PAGE gel, cell lysate and VLP sample volume loading were normalized to sample protein content, determined by a BCA assay. Gels were transferred to a nitrocellulose membrane and immunoblotted was performed as described previously in the section, *Immunoblotting*. Following ECL detection, VP40 cell lysate (VP40_CL_) expression was normalized to the respective GAPDH band density. The relative budding index was calculated according to the ratio of density_VLP_/density_C+VLP_ (where density_VLP_ is the eVP40 VLP band density and density_C+VLP_ is the eVP40 cell lysate + VLP band density).

### Scanning electron microscopy

HEK293 cells were transfected with FLAG-eVP40 and treated as described in the previous section, *Cell Culture, Transfections & Pharmacological Treatments.* Cells were scraped and collected through low-speed centrifugation at 48 hours post-transfection, and stored in primary fixative (2% glutaraldehyde, 2% paraformaldehyde in 0.1 M cacodylate buffer, pH 7.35) at 4°C until processing. During processing, samples were fixed to coverslips and post-stained with 1% osmium tetroxide in 0.1 M cacodylate buffer. Samples were extensively rinsed with water and dehydrated with a graded series of ethanol followed by drying in a Tousimis 931 Supercritical Autosamdri® device. Prior to imaging, samples were coated with 3 nm Iridium. A Field Emission Scanning Electron Microscope Magellan 400 (FEI) (Hillsboro, OR) was used to collect images, with assistance from Tatyana Orlova at the Notre Dame Integrated Imaging Facility.

### Transmission electron microscopy imaging

VLPs were purified as previously described in *VLP Collections & Functional Budding Assays.* Following ultracentrifugation, VLPs were resuspended in fixative (2.5% glutaraldehyde in 0.1 M cacodylate buffer). Purified VLPs were applied onto glow discharged carbon formvar grids and negatively stained using 4% uranyl acetate. Samples were imaged with a FEI Tecnai T12 electron microscope equipped with a tungsten source and operating at 80 kV. VLP length and diameter measurements were quantified using ImageJ software. For diameter analysis, eight different diameters were measured across random areas on each VLP, and the mean diameter was reported.

### Circular dichroism

VLPs were produced in the presence of DMSO or fendiline and purified as previously described in the *VLP collections* section. PB buffer was added to a 10-mm path-length Spectrosil Far UV Quartz cuvette (Starna Cells CatID: 21-Q-10) and a background spectra was collected and autosubtracted from VLP samples. VLP samples were loaded into the 10-mm cuvette diluted in PB buffer to an approximate final protein concentration of 30 µg/mL. Circular dichroism spectra of VLP samples were collected between 200-280 nm at a 0.2 nm step size with 0.5 s time-per-point (with adaptive sampling) using a Chirascan spectrometer (Applied Photophysics, Leatherhead, UK). Absorbance spectra (Abs) and detector signal (hv) were collected simultaneously as controls. After collecting the CD spectra, a microstir bar was added to the cuvette and thermal melting was run from 20°C-93°C using a 0.5°C step ramping at 1.00°C/minute with a tolerance of 0.20°C; simultaneously, spectra were collected at a single wavelength of 220 nm with a time-per-point of 24 seconds. Absorbance spectra (Abs) and detector signal (hv) were collected simultaneously as controls. At the end of thermal melting measurement collection, temperature set-points were replaced with temperatures measured by the sample handling unit. Data was converted from Chirascan filetype to CSV and then extracted into GraphPad PRISM 7. Using the first derivative of the circular dichroism spectral signal in respect to temperature, the maxima was taken as the melting point of the sample. Melting temperatures of the three replicates were averaged for the reported T_m_.

### DiI entry assay

VLPs produced from HEK293 cells expressing FLAG-eVP40 and eGP were purified as previously described in the *Functional Budding Assays* section. DiI entry assays were performed as described previously (Kuroda *et al*, 2015; Nanbo *et al*, 2018). In brief, following ultra-centrifugation VLPs were resuspended in STE buffer and further purified by filtering through a 0.22 µm filter. Protein content of VLP samples were normalized to 0.1 µg/mL using STE buffer. VLPs were labeled with DiI for 1 hr at RT with gentle agitation (final DiI = 0.06 µM). Following incubation, labeled VLP samples were concentrated down to equal volumes, and brought up to volume in phenol-free MEM with 2% FBS and 4% BSA.

HEK293 cells were transfected with TIM-1 for 24 hours prior to incubation with DiI-labeled VLPs and briefly rinsed with phenol-free MEM with 2% FBS and 4%BSA. DiI-VLPs were added to TIM-1 expressing HEK293 cells, spinoculated for 45 min at 4°C, and allowed to incubate for 1 hr at 37°C. Plates were then rinsed with PBS, fixed with 4% paraformaldehyde in PBS, their nuclei stained with Hoechst 3342, and stored at 4°C until imaging. During image acquisition, z-stacks were acquired of 10-15 frames (1 μm steps).

### Toxicity analysis

HEK293 and Vero E6 cell toxicity following fendiline treatment was tested at the indicated time points using the Cell Titer Glo Viability Assay (Promega, Madison WI) according to the manufacturer’s protocol. In brief, HEK293 cells were treated with the indicated concentration of fendiline or control for 24 or 48 hours. Vero E6 cells were treated for 24 hours, the drug was removed and the cells were replenished with Vero E6 culture media, to mirror the corresponding ebolavirus infections at BSL-4. Following the one-hour mock infection, cells were washed with PBS and plates were divided into three treatment groups (day 0, e.d., e.o.d.), and cells received either culture media or freshly prepared fendiline or vehicle control and were then treated daily with freshly prepared compound or left to incubate based on their designated treatment group. At 48, 72, and 96 hours following mock infection, and mirroring the post infection fixation time points, CellTiter-Glo® reagent was added to each well in accordance with the manufacturer’s instructions. Both HEK293 and Vero E6 toxicity assays luminescence readings were recorded using a SpectraMax® M5 (Molecular Devices®) plate reader.

### Mathematical model of *in vitro* experiments

We implemented a system of ordinary differential equations (ODEs) to describe the dynamics of host target cells, infected cells and free virus in different combinations reflecting the *in vitro* experimental systems used here. These equations are similar to those used to simulate Ebola virus dynamics in earlier work (Nguyen *et al*, 2015; Martyushev *et al*, 2016).

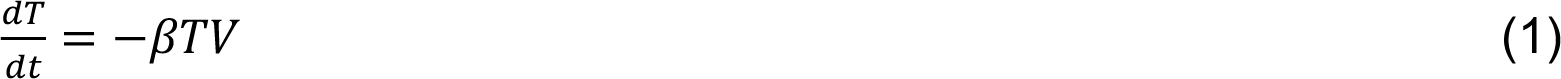

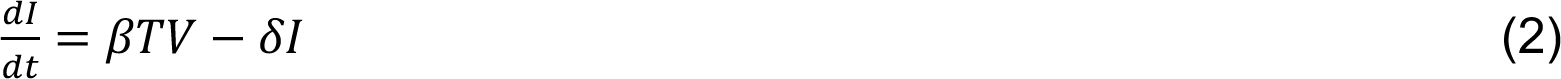

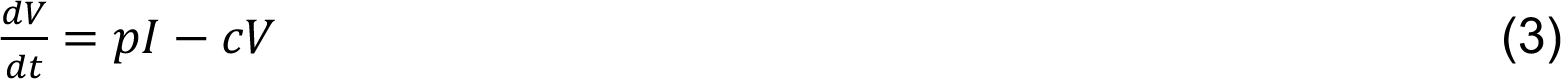

Where *T*, *I* and *V* represent numbers of susceptible target host cells, infected cells and free virions respectively and parameters are described in Table 2.

We modified appropriate parameters in equations 1-3 to represent the following experimental systems during calibration of the mathematical model:

- Viral budding assay (set b=0, T(0) = 0, I(0) = 2.625×10^6^, V(0) = 0, vary d, c, p)
- Viral entry assay (set p=0, T(0) = 6.3×10^5^, I(0) = 0, V(0) = 6.3×10^3^, vary b, d, c)
- BSL-4 Cellular infection assay (BSL4) (T(0) = 5×10^4^, I(0) = 0, V(0) = 5×10^4^ (MOI 1) or V(0) = 5×10^3^ (MOI 0.1), vary b, d, c, p)

Fendiline treatment effects are simulated using E_max_ dose response curves:

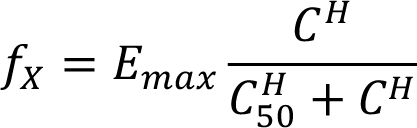

Where *C*: concentration of fendiline, *E_max_*: maximum effect of fendiline, *H*: hill constant for the dose response curve, and *C_50_* concentration with 50% of *E_max_* efficacy. *E_max_*, *C_50_* and *H* are fitted separately for fendiline effects on budding (*f_budding_*) or entry (*f_entry_*). Fendiline efficacy (*f_X_*) is defined as a fraction where *f_X_*=0 implies no effect and *f_X_*=1 implies 100% inhibition of *X* (*X* = budding or entry). Fendiline effects are integrated into equations (1-3) by multiplying b by (1-*f_entry_*) and multiplying p by (1-*f_budding_*). Fendiline concentrations are assumed to be constant over the observation periods based on low *in vitro* degradation rates of the drug. Daily treatment in the BSL-4 assays (e.d.) are simulated by removing all free virus particles from the equations at each dosing time.

We calibrated the model in two stages. First, we calibrated to the budding and entry assays. The uncoupling of budding and entry in this data allows us to define biologically feasible ranges for the effects of fendiline on budding and entry separately. Using these feasible ranges, we proceed to calibrate the full model to the BSL-4 data (day-1/0 and e.d.). Therefore, we allow the budding and entry assays to inform the BSL-4 simulations without imposing strict assumptions about the equivalency between the two systems. In this way we progressively build complexity into the model accounting for fendiline effects on viral budding, entry, and infection progression.

Calibrating to budding and entry assays we restrict the value of p to be larger than 1 and c to be between 0 and 5. These assumptions are in line with previous estimates, and are necessary to qualitatively reproduce viral production observed, but not quantified, in the budding assays. Parameters are estimated using Matlab’s non-linear least squares optimization algorithm. Parameter bounds and final values are defined in Table 2.

### Statistical testing

All experiments were done in triplicate (unless otherwise noted and figure legends indicate the number of replicates). For analysis of eVLP diameter and length from TEM experiments, as well as total PA levels between control and 5 µM fendiline treated cells, a two-tailed t-test was performed. For all experiments which contained >2 experimental groups, a one-way ANOVA with Dunnett’s multiple comparisons was performed on raw data. Lastly, for N&B analysis, a two-way ANOVA with Dunnett’s multiple comparisons was performed.

## Data Availability

No data were deposited in a public database and any additional data from this paper can be requested from the authors.

## Acknowledgements

These studies were supported by the NIH (AI081077) to R.V.S and the Indiana CTSI to E.P. and R.V.S. (UL1TR001108). M.L.H. was partially supported by a NIH T32 fellowship (T32 GM075762), E.A.D. was supported by NIH T32 fellowship (GM132024), and C.B.P. was supported by NIH T32 fellowship (AI148103). We are grateful for lipidomic analysis by Avanti Polar Lipids, Inc., and for research support by Dr. Nathan Dissinger. The authors acknowledge the use of the facilities of the Bindley Bioscience Center, a core facility of the NIH-funded Indiana Clinical and Translational Sciences institute and the use of the Purdue Life Science Electron Microscopy facility.

## Author Contributions

Conceptualization: M.L.H. and R.V.S. Investigation: M.H., S.A., L.I.P., E.A.D., C.B.P., and K.E.H. Methodology: M.H., S.A., L.I.P., E.A.D., C.B.P., K.E.H., and L.I.P. designed the BSL-4 experiments. Formal analysis: M.H., S.A., L.I.P., E.A.D., C.B.P., L.I.P. and K.E.H. performed and analyzed the BSL-4 experiments. E.P. designed and performed the mathematical modeling and analysis. Funding acquisition: M.L.H., E.A.D., C.B.P., E.P., and R.V.S. Project administration: R.V.S. Supervision: R.V.S. oversaw the in vitro and BSL-2 work, J.M.B. and J.D. oversaw the BSL-4 work, E.P. oversaw the mathematical modeling and analysis. Writing - original draft: M.L.H. and R.V.S. Writing – review and editing: all authors.

## Expanded View Figure Legends

**Expanded View 1 (EV1).**
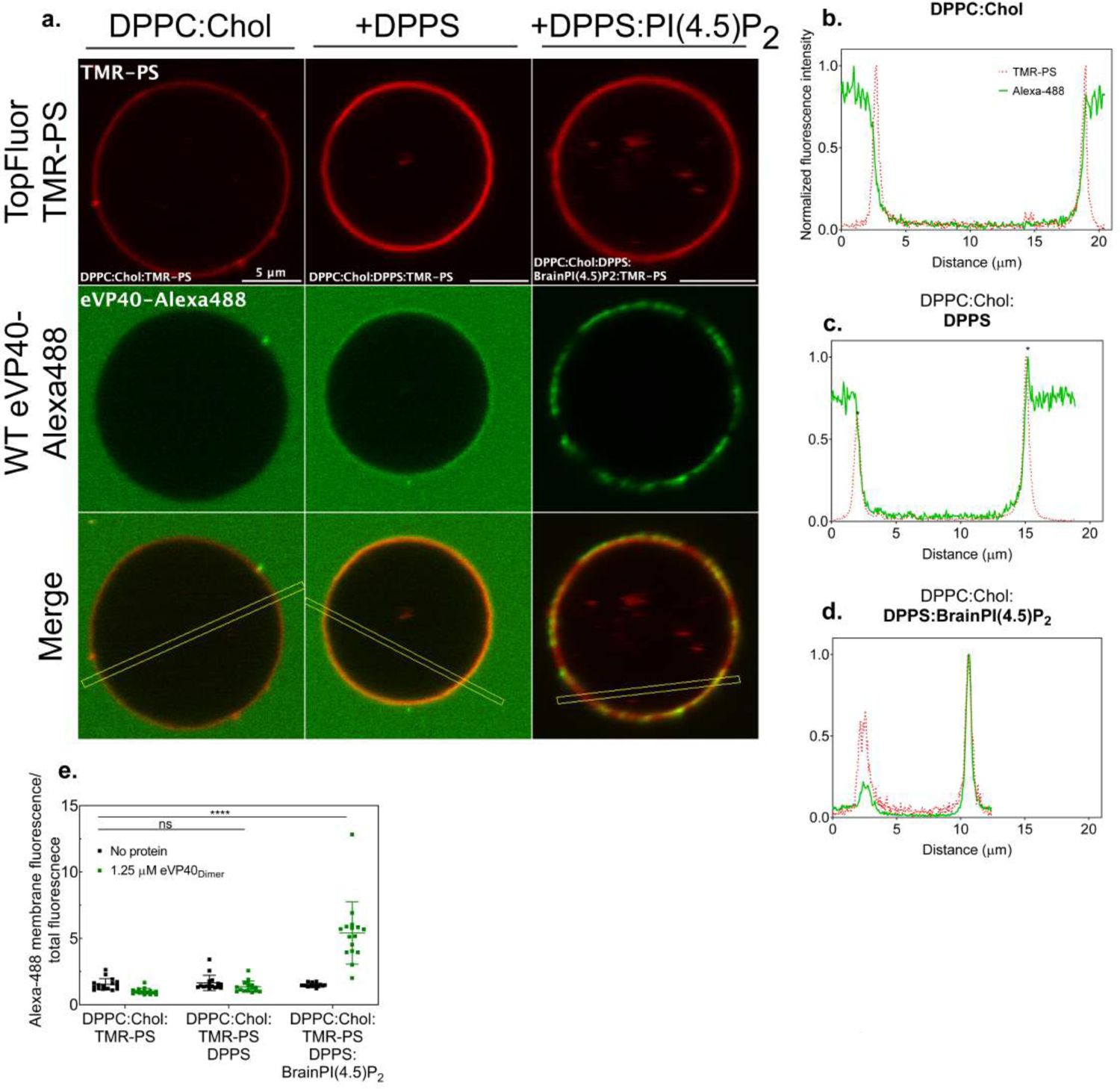
Fluorescence profiles of PS and WT-eVP40 through confocal microscopy and plot profile analysis of fluorescently labeled GUVs and Alexa488-eVP40. **A** Representative confocal images of fluorescently labeled (TopFluor® TMR-PS) GUVs with varying lipid compositions (red) following incubation with eVP40-Alexa88 (green). Colocalization between eVP40 and PS within GUVs was indicated by plot profile analyses of fluorescence signals between TopFluor® TMR-PS (red dotted line) and eVP40- Alexa488 (green solid line) performed at indicated open yellow lines in **a** and shown in **B-D**. **B** Plot profile analysis of eVP40-Alexa488 and control GUVs (DPPC:Cholesterol:0.2mol% TopFluor® TMR-PS), **C** Plot profile analysis of eVP40- Alexa488 and PS GUVs (DPPC:Cholesterol:0.2mol% TopFluor® TMR-PS:DPPS). * indicates overlap in fluorescence signals. **D** Plot profile analysis of eVP40-Alexa488 and PS+PI(4,5)P_2_ GUVs (DPPC:Cholesterol:0.2mol% TopFluor® TMR-PS:DPPS:PI(4,5)P_2_). **E** Plot of WT eVP40 Alexa-488 membrane fluorescence/total fluorescence for lipid compositions shown in a-d, n = 3, ****p<0.0001. PS: phosphatidylserine; DPPC: dipalmitoyl-phosphatidylcholine; DPPS: dipalmitoyl-phosphatidylserine.

**Expanded View 2 (EV2).**
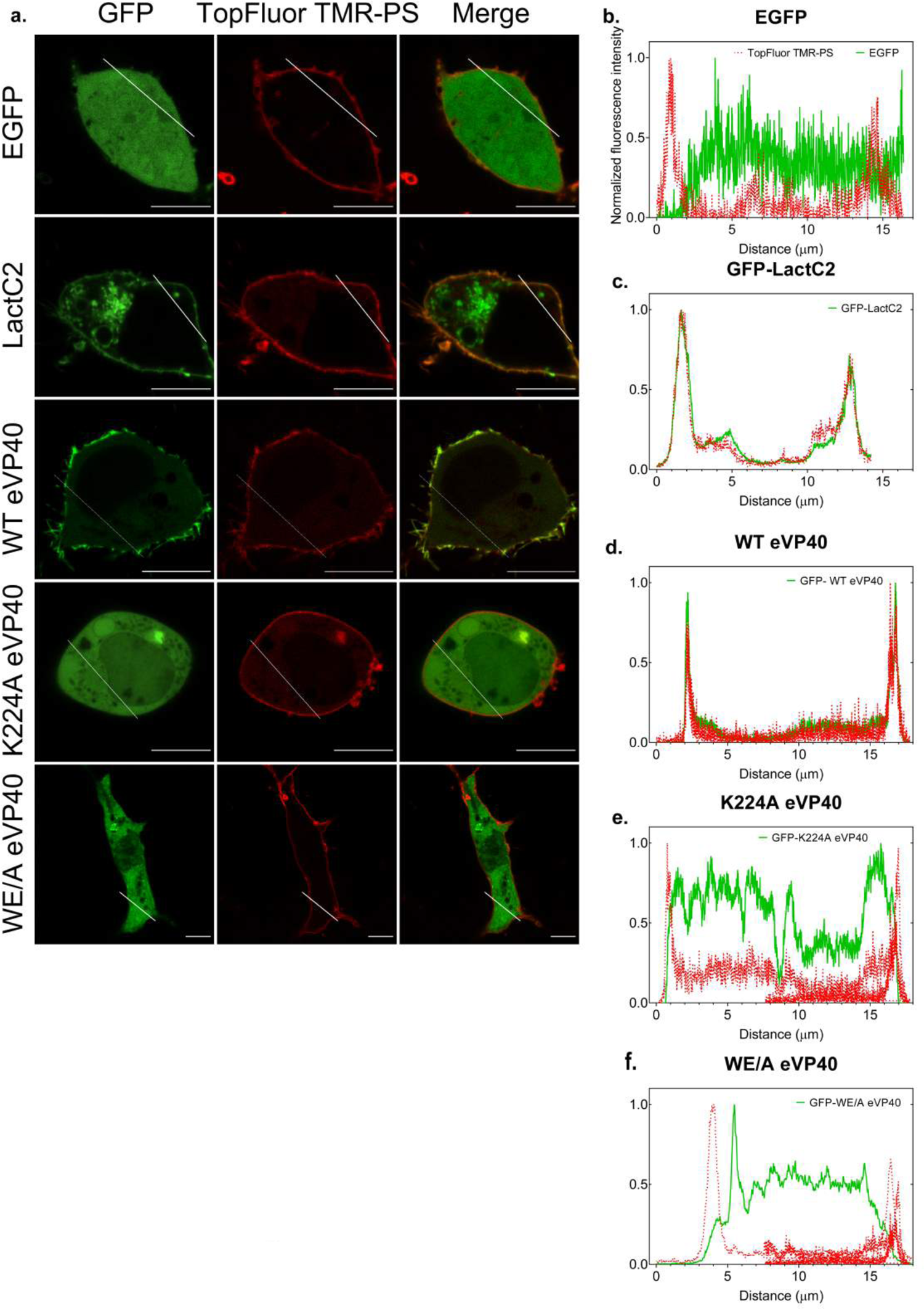
Fluorescence profiles of PS and GFP-WT-eVP40 in HEK 293 cells through confocal microscopy. **A** Representative confocal images from live cell imaging of HEK293 expressing various GFP-fused proteins (GFP; green) following supplementation with TopFluor® TMR-PS (red). Solid white lines indicate where plot profile analysis was performed; scale bar= 10 µm. **B-C** Validation of ability to detect exogenously added fluorescently labelled PS within the inner leaflet of the plasma membrane of cells **B** Plot profile analysis of HEK293 cells expressing cytosolic GFP. **C** Plot profile analysis of HEK293 cells expressing the PS sensor GFP-LactC2 **D-E** Investigation of functionally distinct eVP40 proteins ability to bind to fluorescently labelled PS within the inner leaflet of the plasma membrane in living cells. **D** Plot profile analysis of HEK293 cells expressing GFP-WT-eVP40. **E** Plot profile analysis of HEK293 cells expressing GFP-K224A-eVP40 (PS-binding residue mutant). **F** Plot profile analysis of HEK293 cells expressing GFP- WE/A-eVP40 (oligomerization deficient mutant). TopFluor TMR-PS fluorescence signal intensity (red dotted line) and GFP fluorescence signal intensity (green solid line).

**Expanded View 3 (EV3).**
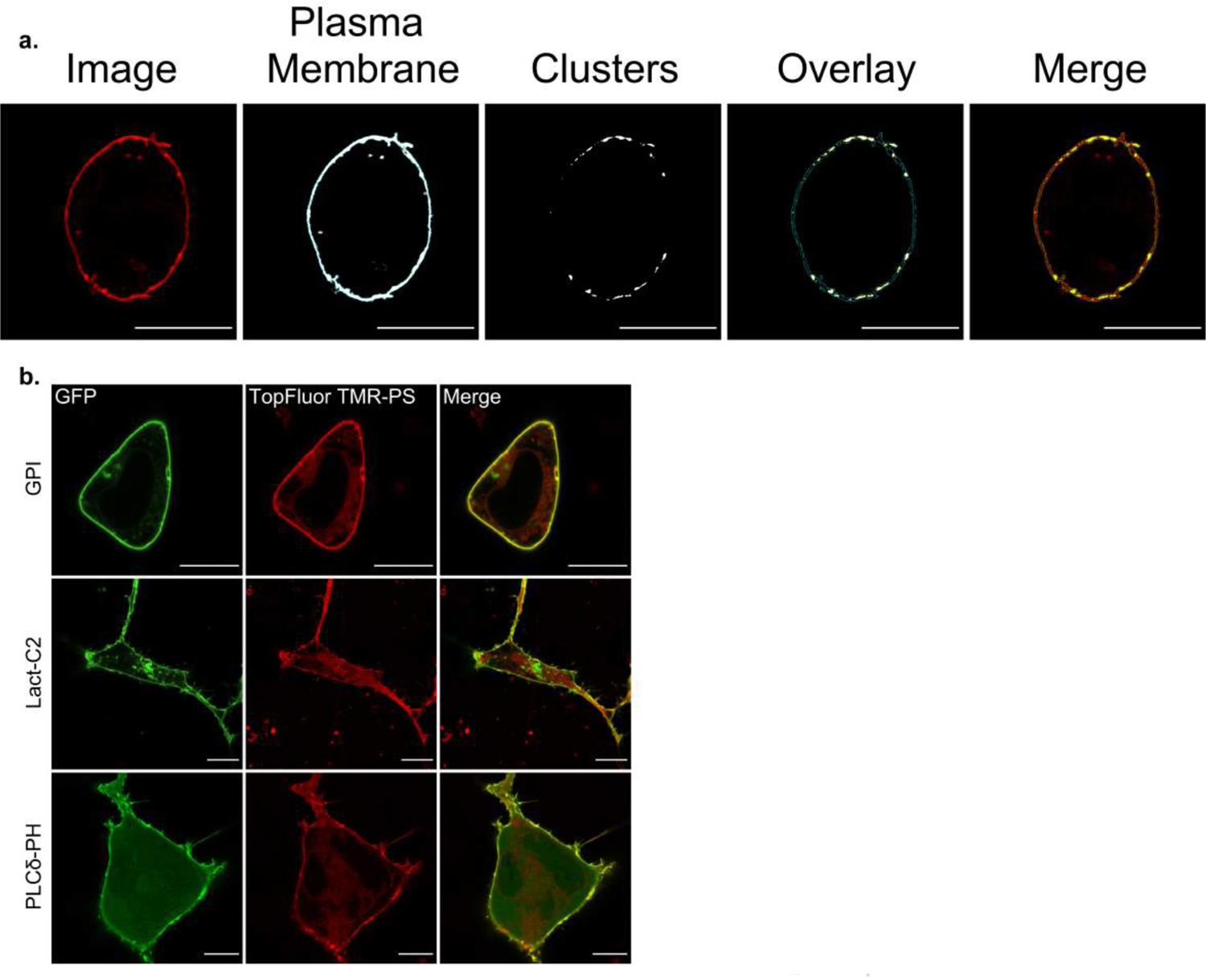
Validation of in vitro and cellular PS clustering experiments. **A** GUVs of varying compositions were imaged prior to and following the addition of 1.25 µM eVP40-Alexa. eVP40-Alexa 488 enrichment ratios at the membrane of the GUVs were calculated by the ratio the Alexa488 fluorescence intensity at the GUV membrane / Alexa488 total fluorescence. Values are reported as mean ± s.d.; of n=3. A two-way ANOVA with multiple comparisons was performed. *p<0.0001. **B** Representative images of the step- wise image analysis workflow of quantifying PS clustering in living HEK293 cells expressing GFP-fused proteins using a custom ImageJ macro. scale bar= 10 µm. **C** Representative images from live cell imaging experiments of HEK293 cells expressing control GFP-fused proteins specific for the plasma membrane (GPI), and specific lipids, PS (LactC2) and PI(4,5)P_2_ (PLCδ-PH). scale bar = 10 µm.

**Expanded View 4 (EV4).**
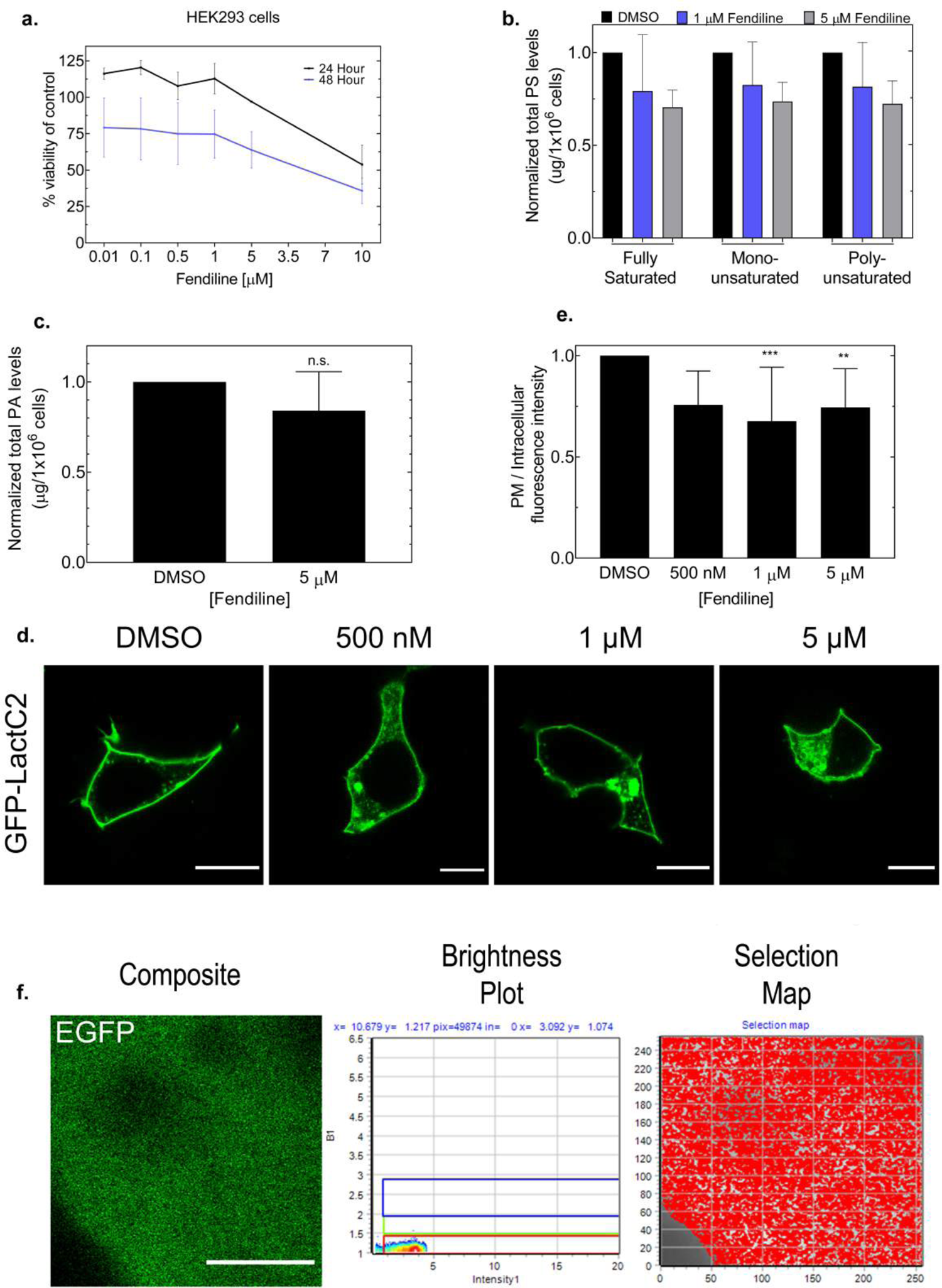
Profile of fendiline toxicity and efficacy in BSL-2 models. **A** CellTiter-Glo® viability results of HEK293 cells. HEK293 cells were treated with fendiline for 24 hours (black line) and 48 hours (blue line) and viability was assessed as a % viability of control (n = 3). **B** PS saturation analysis from lipidomic analysis (LC/MS/MS) of total lipids extracted from HEK293 cells treated with the indicated concentration of fendiline (48 hours). Values are normalized to DMSO control and are reported as mean ± s.d.; n=3; A one-way ANOVA was performed with multiple comparisons compared to the control DMSO. **C** PA level analysis from lipidomic analysis (LC/MS/MS) of total lipids extracted from HEK293 cells treated with 5 µM fendiline (48 hours). Values are normalized to DMSO control and are reported as mean ± s.d.; n=3; A two-tailed t-test was performed. **D-E** Analysis of PS plasma membrane localization in response to 24 hour fendiline treatment. **D** Representative confocal images from live cell imaging of HEK293 cells expressing GFP- LactC2 and treated with fendiline for 24 hours; scale bars = 10 µm. **E** Effect of fendiline on PS plasma membrane localization was calculated by the ratio of GFP fluorescence at the (plasma membrane intensity/intracellular intensity). Values are normalized to DMSO control and are reported as mean ± s.d.; N>15, n=3; A one-way ANOVA was performed with multiple comparisons compared to the DMSO control. **p=0.0045; ***p=0.0003. **f** N&B analysis of HEK293 cells expressing the control GFP. Analysis was performed at 48 hours post treatment (DMSO) to align with N&B analysis performed on experiments with HEK293 cells expressing GFP-LactC2 or GFP-eVP40 and treated with the control or fendiline. *Left panel:* Representative images from time-lapse (30 frames) of HEK293 expressing EGFP and treated with fendiline for 48 hours. scale bar = 5 µm. *Middle panel:* Brightness and Intensity plots for representative image. *Right panel:* Selection map correlating each pixel in the representative image to an oligomerization state (b value) (red: monomer). PS: phosphatidylserine; PA: phosphatidic acid; N&B: number & brightness analysis.

**Expanded View 5 (EV5).**
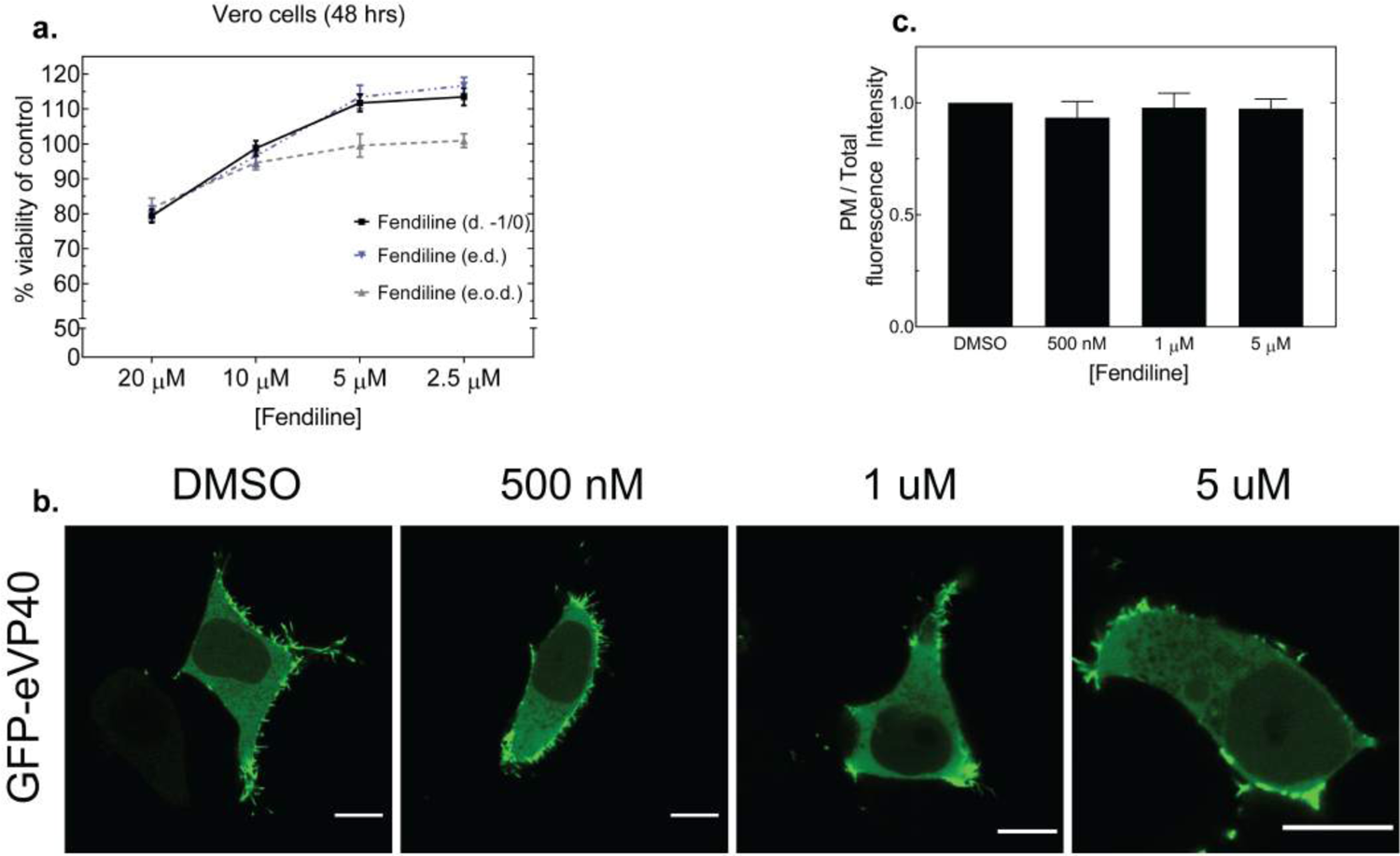
Vero E6 toxicity and effect of fendiline on eVP40 plasma membrane localization at 24 hours post treatment. **A** CellTiter-Glo® viability results of Vero cells. Cells were treated with control or fendiline for 48 hours according to the BSL-4 infection model; d- 1/0 (black line), e.d. (blue line) and e.o.d (gray line) and viability was assessed as a % viability of control (n = 3). **B-C** Effect of fendiline on eVP40 PM localization in HEK293 cells after 24 hours of treatment. **B** Representative confocal images from live cell imaging experiments of HEK293 cells expressing EGFP-WT-eVP40 after 48 hours of fendiline treatment. scale bars= 10 µm. **C** Effect of fendiline on eVP40 PM localization was quantified by the ratio of EGFP fluorescence intensity at the PM / total GFP fluorescence intensity (and normalized to DMSO control). N>15, n=3. Values are reported as mean ± s.d. A one-way ANOVA with multiple comparisons was performed compared to the DMSO control. PM: plasma membrane.

## References

1. Adu-Gyamfi E, Digman MA, Gratton E, Stahelin R V (2012) Investigation of Ebola VP40 assembly and oligomerization in live cells using number and brightness analysis. Biophys J 102: 2517–2525

2. Adu-Gyamfi E, Soni SP, Xue Y, Digman MA, Gratton E, Stahelin RV (2013) The Ebola virus matrix protein penetrates into the plasma membrane: a key step in viral protein 40 (VP40) oligomerization and viral egress. J Biol Chem 288: 5779–5789

3. Adu-Gyamfi E, Johnson KA, Fraser ME, Scott JL, Soni SP, Jones KR, Digman MA, Gratton E, Tessier CR, Stahelin RV (2015) Host Cell Plasma Membrane Phosphatidylserine Regulates the Assembly and Budding of Ebola Virus. J Virol 89: 9440–9453

4. Amara A, Mercer J (2015) Viral apoptotic mimicry. Nat Rev Microbiol 13: 461–469

5. Amiar S, Stahelin RV (2020) The Ebola virus matrix protein VP40 hijacks the host plasma membrane to form virus envelope. J Lipid Res 61: 971

6. Amiar S, Husby ML, Wijesinghe KJ, Angel S, Bhattarai N, Gerstman BS, Chapagain PP, Li S, Stahelin RV (2021) Lipid-specific oligomerization of the Marburg virus protein VP40 is regulated by two distinct interfaces for virion assembly. J Biol Chem 296: 100796

7. Banadyga L, Dolan MA, Ebihara H (2016) Rodent-Adapted Filoviruses and the Molecular Basis of Pathogenesis. J Mol Biol 428: 3449–3466

8. Bayer R, Mannhold R. (1987) Fendiline: a review of its basic pharmacological and clinical properties. Pharmatherapeutica 5: 103–136

9. Bell RM, Ballas LM, Coleman RA (1981) Lipid topogenesis. J Lipid Res 22: 391–403

10. Berjukow S, Doring F, Froschmayr M, Grabner M, Glossman H, Hering S (1996) Endogenous calcium channels in human embryonic kidney (HEK293) cells. Br J Pharmacol 118: 748-753

11. Bobone S, Hilsch M, Storm J, Dunsing V, Herrmann A, Chiantia S (2017) Phosphatidylserine Lateral Organization Influences the Interaction of Influenza Virus Matrix Protein 1 with Lipid Membranes. J Virol 91: 1–15.

12. Bolte S, Cordelières FP (2006) A guided tour into subcellular colocalization analysis in light microscopy. J Microsc 224: 213–232

13. Bornholdt ZA, Noda T, Abelson DM, Halfmann P, Wood MR, Kawaoka Y, Sahpire EO (2013) Structural rearrangement of ebola virus vp40 begets multiple functions in the virus life cycle. Cell 154: 763–77

14. Breman JG, Heymann DL, Lloyd G, McCormick JB, Miatudila M, Murphy FA, Muyembé- Tamfun JJ, Piot P, Ruppol JF, Sureau P, et al (2016) Discovery and Description of Ebola Zaire Virus in 1976 and Relevance to the West African Epidemic during 2013- 2016. J Infect Dis 214: S93–S101

15. Brunton B, Rogers K, Phillips EK, Brouillette RB, Bouls R, Butler NS, Maury W (2019) TIM-1 serves as a receptor for Ebola virus in vivo, enhancing viremia and pathogenesis. PLoS Negl Trop Dis 13: e0006983

16. Carnec X, Meertens L, Dejarnac O, Perera-Lecoin M, Hafirassou ML, Kitaura J, Ramdasi R, Schwartz O, Amara A (2015) The Phosphatidylserine and Phosphatidylethanolamine Receptor CD300a Binds Dengue Virus and Enhances Infection. J Virol 90: 92–102

17. Cheng JS, Wang JL, Lo YK, Chou KJ, Lee KC, Liu CP, Chang HT, Jan CR (2001) Effects of the antianginal drug fendiline on Ca2+ movement in hepatoma cells. Hum Exp Toxicol 20: 359–364

18. Cho KJ, van der Hoeven D, Zhou Y, Maekawa M, Ma X, Chen W, Fairan GD, Hancock JF (2015) Inhibition of Acid Sphingomyelinase Depletes Cellular Phosphatidylserine and Mislocalizes K-Ras from the Plasma Membrane. Mol Cell Biol 36: 363–374

19. Cho W, Stahelin RV (2005) Membrane-protein interactions in cell signaling and membrane trafficking. Annu Rev Biophys Biomol Struct 34: 119–151

20. Darszon A, Vandenberg CA, Schönfeld M, Ellisman MH, Spitzer NC, Montal M (1980) Reassembly of protein-lipid complexes into large bilayer vesicles: perspectives for membrane reconstitution. Proc Natl Acad Sci U S A 77: 239–243

21. Dejarnac O, Hafirassou ML, Chazal M, Versapuech M, Gaillard J, Perera-Lecoin M, Umana-Diaz C, Bonnet-Madin L, Carnec X, Tinevez JY et al (2018) TIM-1 Ubiquitination Mediates Dengue Virus Entry. Cell Rep 23: 1779–1793

22. Del Vecchio K, Frick CT, Gc JB, Oda S, Gerstman BS, Saphire EO, Chapagain PP, Stahelin RV (2018) A cationic, C-terminal patch and structural rearrangements in Ebola virus matrix VP40 protein control its interactions with phosphatidylserine. J Biol Chem 293: 3335–3349

23. Digman MA, Dalal R, Horwitz AF, Gratton E (2008) Mapping the number of molecules and brightness in the laser scanning microscope. Biophys J 94: 2320–2332

24. Dyall J, Nelson EA, DeWald LE, Guha R, Hart BJ, Zhou H, Postnikova E, Logue J, Vargas WM, Gross R, et al (2018) Identification of Combinations of Approved Drugs With Synergistic Activity Against Ebola Virus in Cell Cultures. J Infect Dis 218: S672– S678

25. Fairn GD, Schieber NL, Ariotti N, Murphy S, Kuerschner L, Webb RI, Grinstein S, Parton RG (2011) High-resolution mapping reveals topologically distinct cellular pools of phosphatidylserine. J Cell Biol 194: 257–227

26. Folch J, Lees M, Sloane Stanley GH (1957) A simple method for the isolation and purification of total lipides from animal tissues. J Biol Chem 226: 497–509

27. Graul, AI, Pina, P, Tracy, M, Sorbera, L (2020) This year’s new drugs and biologics. Drugs Today 56: 47–103

28. Han Z, Madara JJ, Herbert A, Prugar LI, Ruthel G, Lu J, Liu Y, Liu W, Liu X, Wrobel JE et al (2015) Calcium regulation of hemorrhagic fever virus budding: Mechanistic implications for host-oriented therapeutic intervention. PLoS Pathog 11: e1005220

29. Harty RN (2018) Hemorrhagic Fever Virus Budding Studies. In: Hemorrhagic Fever Viruses: Methods and Protocols, Salvato MS (ed) pp. 209–215. Springer New York, New York, USA.

30. Hirama T, Das R, Yang Y, Ferguson C, Won A, Yip CM, Kay JG, Grinstein S, Parton RG, Fairn GD (2017) Phosphatidylserine dictates the assembly and dynamics of caveolae in the plasma membrane. J Biol Chem 292: 14292–14307

31. Hoenen T, Biedenkopf N, Zielecki F, Jung S, Groseth A, Feldmann H, Becker S (2010) Oligomerization of Ebola virus VP40 is essential for particle morphogenesis and regulation of viral transcription. J Virol 84: 7053–7063

32. Huang CC, Huang CJ, Cheng JS, Liu SI, Chen IS, Tsai JY, Chou CT, Tseng PL, Jan CR (2009) Fendiline-evoked [Ca2+]i rises and non-Ca2+-triggered cell death in human oral cancer cells. Hum Exp Toxicol 28: 41–48

33. Jan CR, Tseng CJ, Chen WC (2000) Fendiline increases [Ca2+]I in Madin Darby canine kidney (MDCK) cells by releasing internal Ca2+ followed by capacitative Ca2+ entry. Life Sci 66: 1053–1062

34. Jan CR, Lee KC, Chou KJ, Cheng JS, Wang JL, Lo YK, Chang HT, Tang KY, Yu CC, Huang JK (2001) Fendiline, an anti-anginal drug, increases intracellular Ca2+ in PC3 human prostate cancer cells. Cancer Chemother Pharmacol 48: 37–41

35. Jasenosky LD, Cadena C, Mire CE, Borisevich V, Haridas V, Ranjbar S, Nambu A, Bavari S, Soloveva V, Sadukhan S et al (2019) The FDA-Approved Oral Drug Nitazoxanide Amplifies Host Antiviral Responses and Inhibits Ebola Virus. iScience 19: 1279–1290

36. Jasenosky LD, Neumann G, Lukashevich I, Kawaoka Y (2001) Ebola virus VP40-induced particle formation and association with the lipid bilayer. J Virol 75: 5205–5214

37. Jasenosky LD, Kawaoka Y (2004) Filovirus budding. Virus Res. 106: 181–188.

38. Jemielity S, Wang JJ, Chan YK, Ahmed AA, Li W, Monahan S, Bu X, Farzan M, Freeman GJ, Umetsu DT et al (2013) TIM-family Proteins Promote Infection of Multiple Enveloped Viruses through Virion-associated Phosphatidylserine. PLoS Pathog 9: e1003232.

39. Johansen LM, Brannan JM, Delos SE, Shoemaker CJ, Stossel A, Lear C, Hoffstrom BG, Evans Dewald L, Schornberg KL, Scully C et al (2013) FDA-approved selective estrogen receptor modulators inhibit Ebola virus infection. Sci Transl Med 5: 190ra79

40. Johansen LM, DeWald LE, Shoemaker CJ, Hoffstrom BG, Lear-Rooney CM, Stossel A, Nelson E, Delos SE, Simmons JA, Greiner JM et al (2015) A screen of approved drugs and molecular probes identifies therapeutics with anti-Ebola virus activity. Sci Transl Med 7: 290ra89

41. Johnson KA, Budicini MR, Urata S, Gerstman BS, Chapagain PP, Li S, Stahelin RV (2018) PI(4,5)P2 Binding Sites in the Ebola Virus Matrix Protein Modulate Assembly and Budding. bioRxiv doi:10.1101/341248

42. Johnson, KA, Bhattarai N, Budicini MR, LaBonia CM, Baker SCB, Gerstman BS, Chapagain PP, Stahelin RV (2021) Cysteine mutations in the Ebolavirus matrix protein VP40 promote Phosphatidylserine bniding by increasing the flexibility of a lipid-binding loop. Viruses 13: 1375

43. Johnson KA, Taghon GJF, Scott JL, Stahelin R V (2016) The Ebola Virus matrix protein, VP40, requires phosphatidylinositol 4,5-bisphosphate (PI(4,5)P2) for extensive oligomerization at the plasma membrane and viral egress. Sci Rep 6: 19125

44. Kay JG, Koivusalo M, Ma X, Wohland T, Grinstein S (2012) Phosphatidylserine dynamics in cellular membranes. Mol Biol Cell 23: 2198–2212

45. Kerr D, Tietjen GT, Gong Z, Tajkhorshid E, Adams EJ, Lee KYC (2018) Sensitivity of peripheral membrane proteins to the membrane context: A case study of phosphatidylserine and the TIM proteins. Biochim Biophys acta Biomembr 1860: 2126–2133

46. Kondratowicz AS, Lennemann NJ, Sinn PL, Davey RA, Hunt CL, Moller-Tank S, Meyerholz DK, Rennert P, Mullins PF, Brindley M, et al (2011) T-cell immunoglobulin and mucin domain 1 (TIM-1) is a receptor for Zaire Ebolavirus and Lake Victoria Marburgvirus. Proc Natl Acad Sci U S A 108: 8426–8431

47. Kuroda M, Fujikura D, Nanbo A, Marzi A, Noyori O, Kajihara M, et al (2011) Interaction Between TIM-1 and NPC1 Is Important for Cellular Entry of Ebola Virus. J Virol 89: 6481–6493

48. Kurovetz WR, Brunner F, Beubler E, Weyhenmeyer R, Lohaus R, Grob M, Mayer D (1982) Single dose pharmacokinetics of fendiline in humans. Eur J Drug Metab Pharmacokinet 7: 105–110

49. Licata JM, Simpson-Holley M, Wright NT, Han Z, Paragas J, Harty RN (2003) Overlapping Motifs (PTAP and PPEY) within the Ebola Virus VP40 Protein Function Independently as Late Budding Domains: Involvement of Host Proteins TSG101 and VPS-4. J Virol 77: 1812–1819

50. Liu L (2014) Fields Virology, 6th Edition. Clin Infect Dis 59: 613–613

51. Llorente A, Skotland T, Sylvänne T, Kauhanen D, Rog T, Orlowski A, Vattulainen I, Ekroos K, Sandvig K (2013) Biochim Biophys Acta 1831: 1302-1309

52. Madara JJ, Han Z, Ruthel G, Freedman BD, Harty RN (2015) The multifunctional Ebola virus VP40 matrix protein is a promising therapeutic target. Future Virol 10: 537–546.

53. Maekawa M, Fairn GD (2015) Complementary probes reveal that phosphatidylserine is required for the proper transbilayer distribution of cholesterol. J Cell Sci 128: 1422–1433.

54. Maekawa M, Lee M, Wei K, Ridgway ND, Fairn GD (2016) Staurosporines decrease ORMDL proteins and enhance sphingomyelin synthesis resulting in depletion of plasmalemmal phosphatidylserine. Sci Rep 6: 35762

55. Martyushev A, Nakaoka S, Sato K, Noda T, Iwami S (2016) Modelling Ebola virus dynamics: Implications for therapy. Antiviral Res 135: 62–73.

56. Menke M, Gerke V, Steinem C (2005) Phosphatidylserine membrane domain clustering induced by annexin A2/S100A10 heterotetramer. Biochemistry 44: 15296–15303

57. Miller ME, Adhikary S, Kolokoltsov AA, Davey RA (2012) Ebolavirus Requires Acid Sphingomyelinase Activity and Plasma Membrane Sphingomyelin for Infection. J Virol 86: 7473–7483

58. Moe JB, Lambert RD, Lupton HW (1981) Plaque assay for Ebola virus. J Clin Microbiol 13: 791–793.

59. Moller-Tank S, Kondratowicz AS, Davey RA, Rennert PD, Maury W (2013) Role of the phosphatidylserine receptor TIM-1 in enveloped-virus entry. J Virol 87: 8327–8341.

60. Moller-Tank S, Maury W (2014) Phosphatidylserine receptors: Enhancers of enveloped virus entry and infection. Virology 468–470: 565–580.

61. Mühlberger E (2007) Filovirus replication and transcription. Future Virol 2: 205–215.

62. Mullard, A (2020) FDA approves antibody cocktail for Ebola virus. Nat Rev Drug Discov 19: 827

63. Nanbo A, Imai M, Watanabe S, Noda T, Takahashi K, Neumann G, Halfmann P, Kawoaka Y (2010) Ebolavirus is internalized into host cells via macropinocytosis in a viral glycoprotein-dependent manner. PLoS Pathog 6: e1001121

64. Nanbo A, Kawaoka Y (2019) Molecular Mechanism of Externalization of Phosphatidylserine on the Surface of Ebola Virus Particles. DNA Cell Biol 38: 115– 120

65. Nanbo A, Maruyama J, Imai M, Ujie M, Fujioka Y, Nishide S, Takada A, Ohba Y, Kawoaka Y (2018) Ebola virus requires a host scramblase for externalization of phosphatidylserine on the surface of viral particles. PLoS Pathog 14: e1006848

66. Nelson EA, Barnes AB, Wiehle RD, Fontenot GK, Hoenen T, White JM (2016) Clomiphene and Its Isomers Block Ebola Virus Particle Entry and Infection with Similar Potency: Potential Therapeutic Implications. Viruses 8: 206.

67. Nguyen VK, Binder SC, Boianelli A, Meyer-Hermann M, Hernandez-Vargas EA (2015) Ebola virus infection modeling and identifiability problems. Front Microbiol. 6: 257

68. Noda T, Sagara H, Suzuki E, Takada A, Kida H, Kawaoka Y (2002) Ebola Virus VP40 Drives the Formation of Virus-Like Filamentous Particles Along with GP. J Virol 76: 4855–4865

69. Oda S, Noda T, Wijesinghe KJ, Halfmann P, Bornholdt ZA, Abelson DM, Armbrust T, Kawoaka Y, Stahelin RV, Saphire EO (2015) Crystal Structure of Marburg Virus VP40 Reveals a Broad, Basic Patch for Matrix Assembly and a Requirement of the N-Terminal Domain for Immunosuppression. J Virol 90: 1839–1848

70. Panchal RG, Ruthel G, Kenny TA, Kallstrom GH, Lane D, Badie SS, Li L, Bavari S, Aman MJ. (2003) In vivo oligomerization and raft localization of Ebola virus protein VP40 during vesicular budding. Proc Natl Acad Sci U S A 100: 15936–15941

71. Reeves JP, Dowben RM (1969) Formation and properties of thin-walled phospholipid vesicles. J Cell Physiol 73: 49–60

72. Reimão JQ, Mesquita JT, Ferreira DD, Tempone AG (2016) Investigation of calcium channel blockers as antiprotozoal agents and their interference in the metabolism of Leishmania (L.) infantum. Evid Based Complement Alternat Med 2016: 1523691

73. Ruigrok RW, Schoehn G, Dessen A, Forest E, Volchkov V, Dolnik O, Klenk HD, Weissenhorn W (2000) Structural characterization and membrane binding properties of the matrix protein VP40 of Ebola virus. J Mol Biol 300: 103–112

74. Scianimanico S, Schoehn G, Timmins J, Ruigrok RH, Klenk HD, Weissenhorn W (2000) Membrane association induces a conformational change in the Ebola virus matrix protein. EMBO J 19: 6732–6741

75. Skotland T, Sandvig K (2019) The role of PS 18:0/18:1 in membrane function. Nat Commun 10: 2752

76. Soni SP, Stahelin RV (2014) The Ebola virus matrix protein VP40 selectively induces vesiculation from phosphatidylserine-enriched membranes. J Biol Chem 289: 33590–33597

77. Stahelin RV (2014) Membrane binding and bending in Ebola VP40 assembly and egress. Front Microbiol 5: 300

78. Stahelin RV, Scott JL, Frick CT (2014) Cellular and molecular interactions of phosphoinositides and peripheral proteins. Chem Phys Lipids 182: 3–18.

79. Tsai WH (1985) Moment-preserving thresolding: A new approach. *Comput Vision*, Graph Image Process 29: 377–393

80. van der Hoeven D, Cho K, Ma X, Chigurupati S, Parton RG, Hancock JF (2013) Fendiline inhibits K-Ras plasma membrane localization and blocks K-Ras signal transmission. Mol Cell Biol 33: 237–251

81. van der Hoeven D, Cho K, Zhou Y, Ma X, Chen W, Naji A, Montufar-Solis D, Zuo Y, Kovar SE, Levental I et al (2018) Sphingomyelin Metabolism Is a Regulator of K-Ras Function. Mol Cell Biol 38: e00373–17

82. Van Meer G, Voelker DR, Feigenson GW (2008) Membrane lipids: Where they are and how they behave. Nat Rev Mol Cell Biol 9: 112–124.

83. Wan W, Clarke M, Norris MJ, Kolesnikova L, Koehler A, Bornholdt ZA, Becker S, Saphire EO, Briggs JA (2020) Ebola and Marburg virus matrix layers are locally ordered assemblies of VP40 dimers. Elife 9: e592225

84. Wang YE, Park A, Lake M, Pentecost M, Torres B, Yun TE, Wolf MC, Holbrook MR, Freiberg AN, Lee B (2010) Ubiquitin-regulated nuclear-cytoplasmic trafficking of the Nipah virus matrix protein is important for viral budding. PLoS Pathog 6: e1001186

85. Wang J, Qiao L, Hou Z, Luo G (2017) TIM-1 Promotes Hepatitis C Virus Cell Attachment and Infection. J Virol 91: e01583–16

86. Wang P, van der Hoeven D, Ye N, Chen H, Liu Z, Ma X, Montufar-Solis D, Rehl KM, Cho KJ, Thapa S et al (2021) Scaffold repurposing of fendiline: Identification of potent KRAS plasma membrane localization inhibitors. Eur J Med Chem 217: 113381.

87. Watt A, Moukambi F, Banadyga L, Groseth A, Callison J, Herwig A, Ebihara H, Feldmann H, Hoenen T (2014) A Novel Life Cycle Modeling System for Ebola Virus Shows a Genome Length-Dependent Role of VP24 in Virus Infectivity. J Virol 88: 10511– 10524

88. Wen Y, Vogt VM, Feigenson GW (2018) Multivalent Cation-Bridged PI(4,5)P2 Clusters Form at Very Low Concentrations. Biophys J 114: 2630–2639

89. Weyhenmeyer R, Fenzl E, Apecechea M, Rehm KD, Dyde CJ, Johnson KJ, Friedel R (1987) Tolerance and pharmacokinetics of oral fendiline. Arzneimittelforschung 37: 58–62

90. Wijesinghe KJ, Stahelin V (2015) Investigation of the Lipid Binding Properties of the Marburg Virus. J Virol 90: 3074–3085

91. Xiao X, Wang C, Chang D, Wang Y, Dong X, Jiao T, Zhao Z, Ren L, Dela Cruz CS, Sharma L et al (2020) Identification of potent and safe antiviral therapeutic candidates against SARS-CoV-2. Front Immunol 11: 586572

92. Yamashita Y, Oka M, Tanaka T, Yamazaki M (2002) A new method for the preparation of giant liposomes in high salt concentrations and growth of protein microcrystals in them. Biochim Biophys Acta 1561: 129–134

93. Yeung T, Gilbert GE, Shi J, Silvius J, Kapus A, Grinstein S (2008) Membrane phosphatidylserine regulates surface charge and protein localization. Science 319: 210–213

94. Younan P, Iampietro M, Santos RI, Ramanathan P, Popov VL, Bukreyev A (2018) Disruption of Phosphatidylserine Synthesis or Trafficking Reduces Infectivity of Ebola Virus. J Infect Dis 218: S475–S485

95. Zhao H, Hakala M, Lappalainen P (2010) ADF/cofilin binds phosphoinositides in a multivalent manner to act as a PIP(2)-density sensor. Biophys J 98: 2327–2336

