## Appendix (Supplemental data) for "The Ebola virus matrix protein clusters phosphatidylserine, a critical step in viral budding"

**Table of Contents** **Page Number**

Appendix Figure S1 2-3

Thermal melting of eVLPs produced in the presence and absence of fendiline.

Appendix Figure S2 4

Calibration results from the first and second phase of model development.

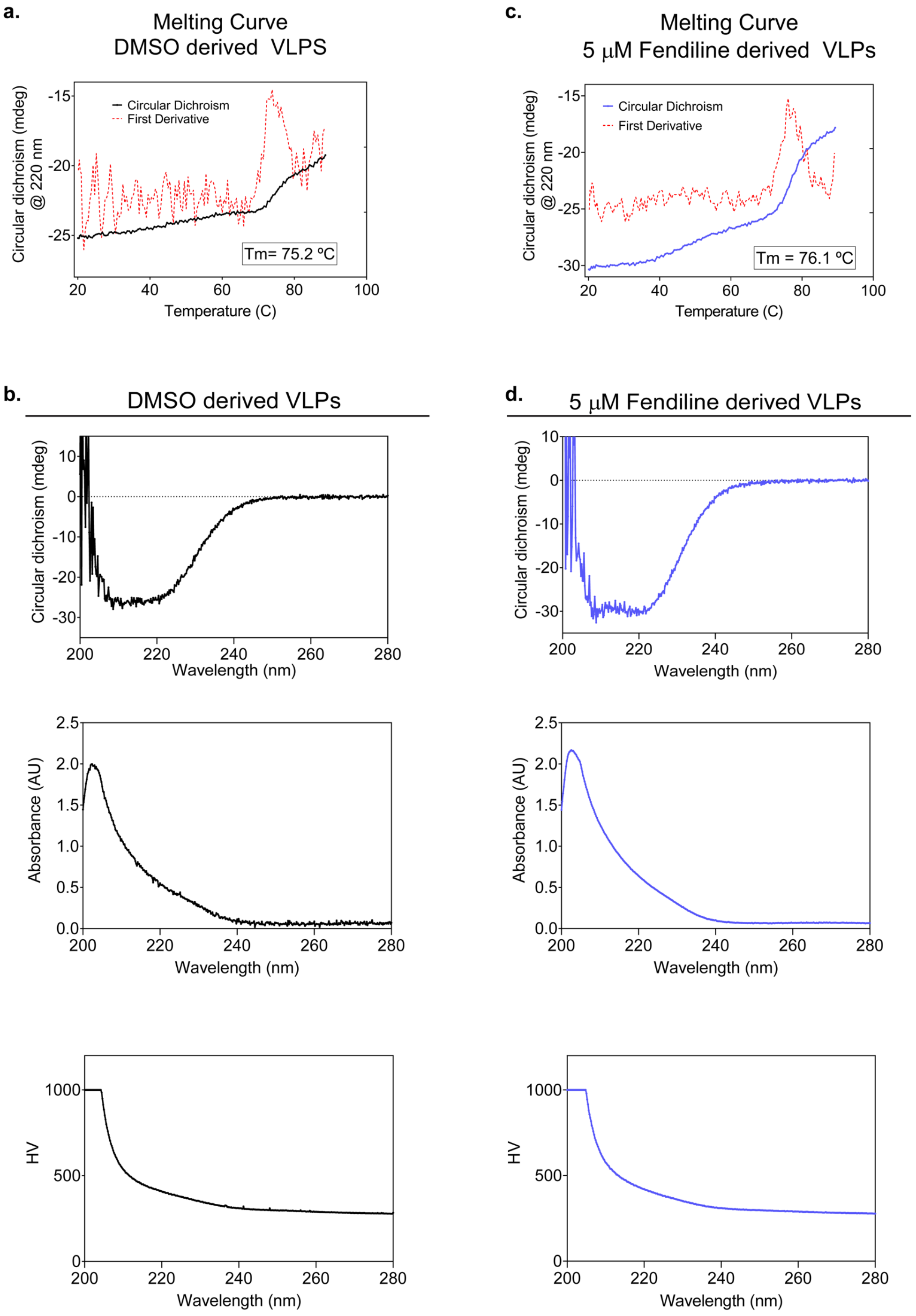


### Appendix Figure S1

**Thermal melting of eVLPs produced in the presence and absence of fendiline. A**

Ebola virus-like particles (eVLPs) were produced in DMSO treated HEK293 cells expressing FLAG-eVP40 and eGP constructs which were then collected, purified, and melted while measuring the loss in polarized light absorption at 220 nm. **B** Prior to melting, the circular dichroism spectra of DMSO VLPs was measured alongside the absorbance and detector (hv) signals. **C** Ebola VLPs were produced in 5 μM fendiline treated HEK293 cells using the same FLAG-eVP40 and eGP constructs which were then collected, purified and melted while measuring the loss in polarized light absorption at 220 nm. **D** Prior to melting, the circular dichroism spectra of fendiline VLPs was measured alongside the absorbance and detector (hv) signals.

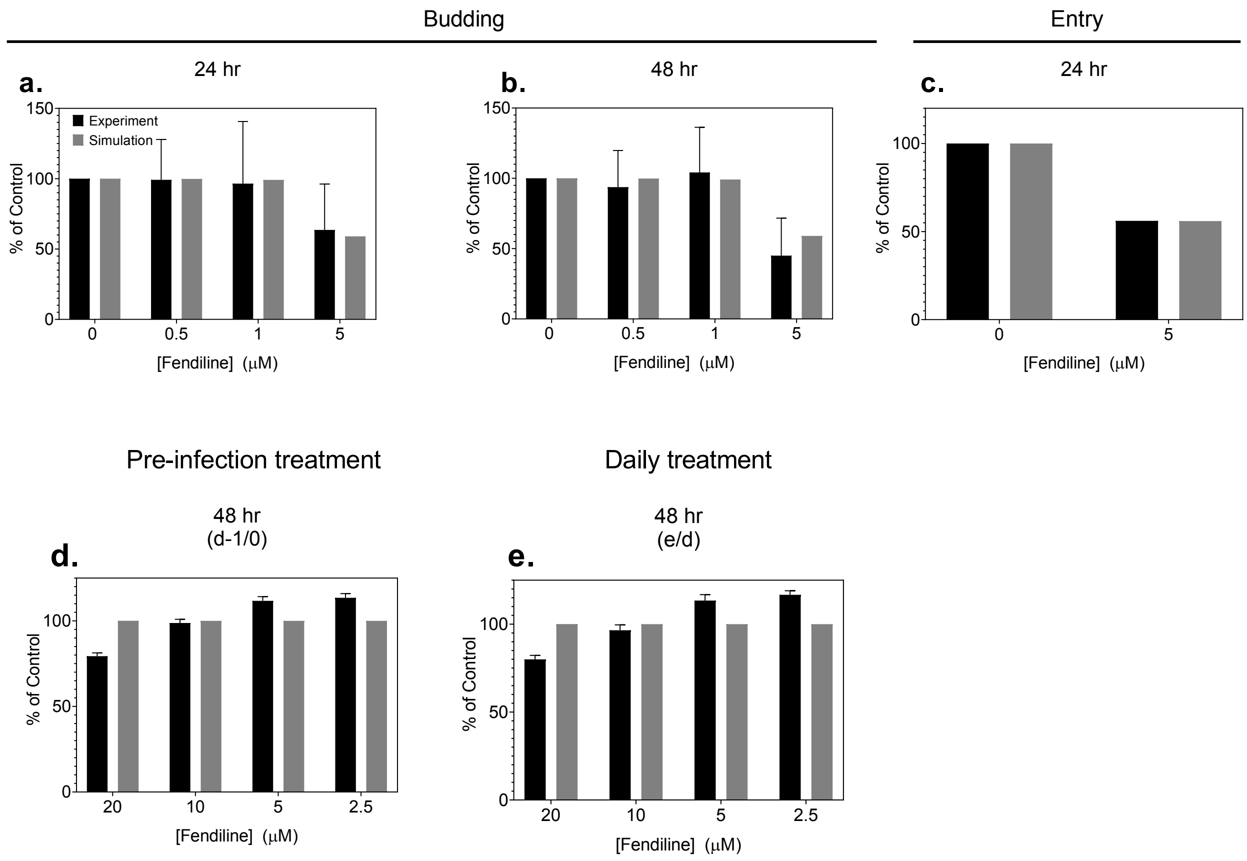


**Appendix Figure S2**

**Calibration results from the first and second phase of model development. A-C** First phase calibration results between experimental (black bars) and simulation (gray) data from budding **(A-B)** and entry **(C)** assays. **D-E** Second phase calibration results showing comparison between experimental and simulation data from cell viability assays. The mathematical model was calibrated to this data and the data in Figure 8 in the main text simultaneously.
